# MOFTy: Multimodal Gaussian Process Factor Analysis with Numerical Information Field Theory

**DOI:** 10.64898/2026.08.12.744240

**Authors:** Maximilian Neumann, Philipp Arras, Anne-Kristin Kaster, Andreas Ott

**Affiliations:** Institute for Mathematics, Heidelberg University, Mathematikon, Im Neuenheimer Feld 205, 69120 Heidelberg, Germany; Institute for Biological Interfaces 5 (IBG-5), Biotechnology and Microbial Genetics, Karlsruhe Institute of Technology (KIT), Hermann-von-Helmholtz-Platz 1, 76344 Eggenstein-Leopoldshafen, Germany; No affiliation, Eisenhartstraße 19, 81245 München, Germany; Institute of Applied Biosciences (IAB), Karlsruhe Institute of Technology (KIT), Fritz-Haber-Weg 4, 76131 Karlsruhe, Germany

## Abstract

Multimodal Gaussian process factor analysis provides a flexible framework for dimensionality reduction in temporally or spatially resolved omics data. Existing approaches, however, typically rely on pre-specified Gaussian process kernel families and do not explicitly separate each latent factor into a component capturing gradual, smooth variation and a complementary component capturing fine-scale, non-smooth variation. Here, we present MOFTy, a Bayesian multimodal factor analysis framework based on numerical information field theory (NIFTy) that replaces fixed kernel families with the flexible correlated field model in NIFTy and enables explicit additive component separation within each latent factor with quantified uncertainty. NIFTy has been successfully applied to high-resolution Bayesian imaging in astrophysics and facilitates scalable, curvature-aware variational inference for efficient posterior approximations. We validate MOFTy on simulated data; applications to published multi-omics data demonstrate that MOFTy disentangles latent spatial structures by separating smooth gradients from localized fine-scale heterogeneity in human glioblastoma and recovers cross-modal patterns in a mouse gastrulation dataset.

## 1 Introduction

Advances in high-throughput sequencing and molecular profiling are generating increasingly complex multimodal datasets with rich spatial and temporal structure [1–4]. Dimensionality reduction is a key strategy for analyzing such data, summarizing high-dimensional relationships in lower-dimensional latent spaces. While deep learning approaches [5–8] offer substantial expressive power, linear matrix factorization models remain widely used due to their interpretability [9–17].

For multi-omics data integration, MOFA [11] introduced a Bayesian factor analysis framework for modeling variation that is shared across or specific to individual omic modalities. It was subsequently extended to multi-group settings [12] and further enhanced in MEFISTO [13] to model dependencies among samples along continuous covariates (e.g., time or space) by placing Gaussian process (GP) priors over the latent factors. More recently, MOFA-FLEX [14] unified and extended these frameworks by adding support for non-negativity constraints, guided factor learning, and the incorporation of prior domain knowledge. Despite these advances, several challenges remain in how these frameworks model covariate-dependent structure with GP priors, which we discuss here for multimodal single-group settings.

First, the correlation structure in each factor is modeled using a stationary and isotropic parametric kernel, either a radial basis function (RBF) or a Matérn kernel. This leaves a single correlation length per factor to be learned from the data, so that variation at several characteristic scales cannot be represented simultaneously.

Second, these parametric kernels are augmented with an additive uncorrelated term, but the two contributions are not inferred as distinct latent fields. Instead, their relative weight is summarized by a factor-specific variance ratio, and component-wise uncertainty estimates are not provided. Inferring the two contributions as distinct latent fields, with uncertainty estimates for each component, could therefore provide a more detailed characterization of each latent factor by disentangling gradual, large-scale structure from fine-scale variation.

Third, these frameworks employ variational inference in which the approximate posterior factorizes across model variables that are not governed by GP priors. Dependencies across these factorization boundaries are therefore not represented. For scalability, sparse inducing-point approximations [18, 19] may additionally be employed for the GP-distributed factors, restricting the approximate posterior covariance to a low-rank structure determined by the number and placement of inducing points. In addition, the GP kernel parameters governing the correlation structure are point-estimated, so their uncertainty does not propagate to the factors.

Here, we introduce MOFTy (see https://github.com/neumann-mn/mofty), a Bayesian multimodal factor analysis framework for single-group datasets with continuous covariates that addresses these challenges using numerical information field theory (NIFTy [20–25]).

MOFTy replaces fixed GP kernel families with the correlated field model (CFM) in NIFTy. The CFM assumes a homogeneous and isotropic correlation structure, which admits a diagonal covariance representation in harmonic space by the Wiener–Khinchin theorem [26, 27] and is fully characterized by its power spectrum. Rather than fixing the spectral shape through a parametric kernel, the CFM models the power spectrum of each factor nonparametrically. Field and power spectrum are inferred jointly, so that uncertainty in the correlation structure propagates to the factors. Scalar spectral hyperparameters are point-estimated, whereas the spectral shape itself is inferred with full posterior uncertainty. The diagonal representation additionally avoids explicit construction of dense covariance matrices, so that memory consumption scales linearly with the number of grid points.

A key feature of MOFTy is the explicit additive decomposition of each latent factor into a CFM component and a complementary non-CFM component, unconstrained by the correlated field prior. Both components are treated as inference targets, so that variational posterior samples provide approximate uncertainty estimates for each component field. As an illustrative example, in spatial dual-omics data from human glioblastoma, this decomposition separates, within each latent factor, gradual, smooth spatial patterns (CFM component) from fine-scale variation and localized heterogeneity (non-CFM component).

For inference, MOFTy employs geometric variational inference (geoVI [28]), a nonlinear extension of metric Gaussian variational inference (MGVI [29]). In geoVI, inference is performed on a Riemannian manifold defined by the Bayesian Fisher information metric of the joint distribution, which encodes local curvature and approximates the posterior precision; MGVI uses the same metric, but only at the current mean. In settings where the posterior has a curved geometry, a Gaussian approximation in standardized coordinates may not be adequate. To address this challenge, geoVI constructs a nonlinear coordinate transformation that makes the metric approximately Euclidean in a neighborhood of an expansion point. A zero-mean, unit-variance Gaussian approximation in these transformed coordinates can then capture the local curvature of the posterior in the original coordinates. In MOFTy, latent variables enter the model multiplicatively at several levels—between the latent factors and the view-specific weights, and within the correlated field model. These couplings, together with the heavy-tailed shrinkage priors on the weights, lead to a curved posterior geometry, which makes geoVI an appropriate choice for MOFTy. MGVI and geoVI are implemented in NIFTy and use an implicit representation of the metric, enabling scalable inference in high-dimensional latent spaces. For large-scale datasets, GPU-accelerated optimization is also supported.

## 2 Results

MOFTy is designed for the analysis of multimodal omics datasets in which features (e.g., genes, proteins, or metabolites) are organized into different views corresponding to molecular modalities (e.g., transcriptomics, proteomics, or metabolomics) and are measured on a shared set of samples, which may correspond to patients, single cells, or spatial spots, with observations allowed to be missing in individual views. The full model specification, together with a discussion of its relation to existing frameworks, is provided in Methods (Section 4).

To begin with, we provide a brief overview of the main model components. The input consists of preprocessed data matrices *Y* ^(1)^, …, *Y* ^(*M*)^ (views), where each matrix *Y* ^(*m*)^ consists of *D*_*m*_ features (rows) and *N* samples (columns) shared across views. As a Bayesian probabilistic model, MOFTy naturally accommodates missing observations across views without requiring prior imputation. Like previous multi-omics factor analysis frameworks [11–14], MOFTy supports Gaussian, Bernoulli, and Poisson observation models—typically used for real-valued, binary, and count data, respectively—so that heterogeneous data types can be modeled jointly, and places heavy-tailed shrinkage priors on the factor weights to improve interpretability of factor-feature associations. For a given number of latent factors *L*, each data matrix *Y* ^(*m*)^ is represented by a low-rank factorization of the form *W* ^(*m*)^*Z*, where *Z* is the *L×N* latent factor matrix shared across all views and *W* ^(*m*)^ is the view-specific *D*_*m*_ *× L* weight matrix. For views with Gaussian likelihoods, the observation model is

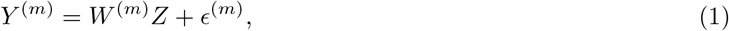

where *ϵ*^(*m*)^ is an additive Gaussian noise term with feature-specific variance inferred from the data. If each sample is additionally associated with a position along a continuous covariate, such as time or space, MOFTy models covariate-dependent variation using the correlated field model (CFM) implemented in NIFTy [20–25]. MOFTy then infers the additive decomposition

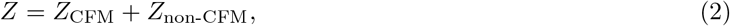

where each row of *Z*, and hence each latent factor, is decomposed into a CFM component and a complementary non-CFM component that captures variation not explained by the correlated field model. Fig. 1a illustrates this decomposition for a factorization along a one-dimensional covariate. The final matrix factorization is reported as the posterior mean together with uncertainty estimates for the factor weights, the combined factors, their CFM and non-CFM components, and the underlying correlation structure.

**Fig. 1.**
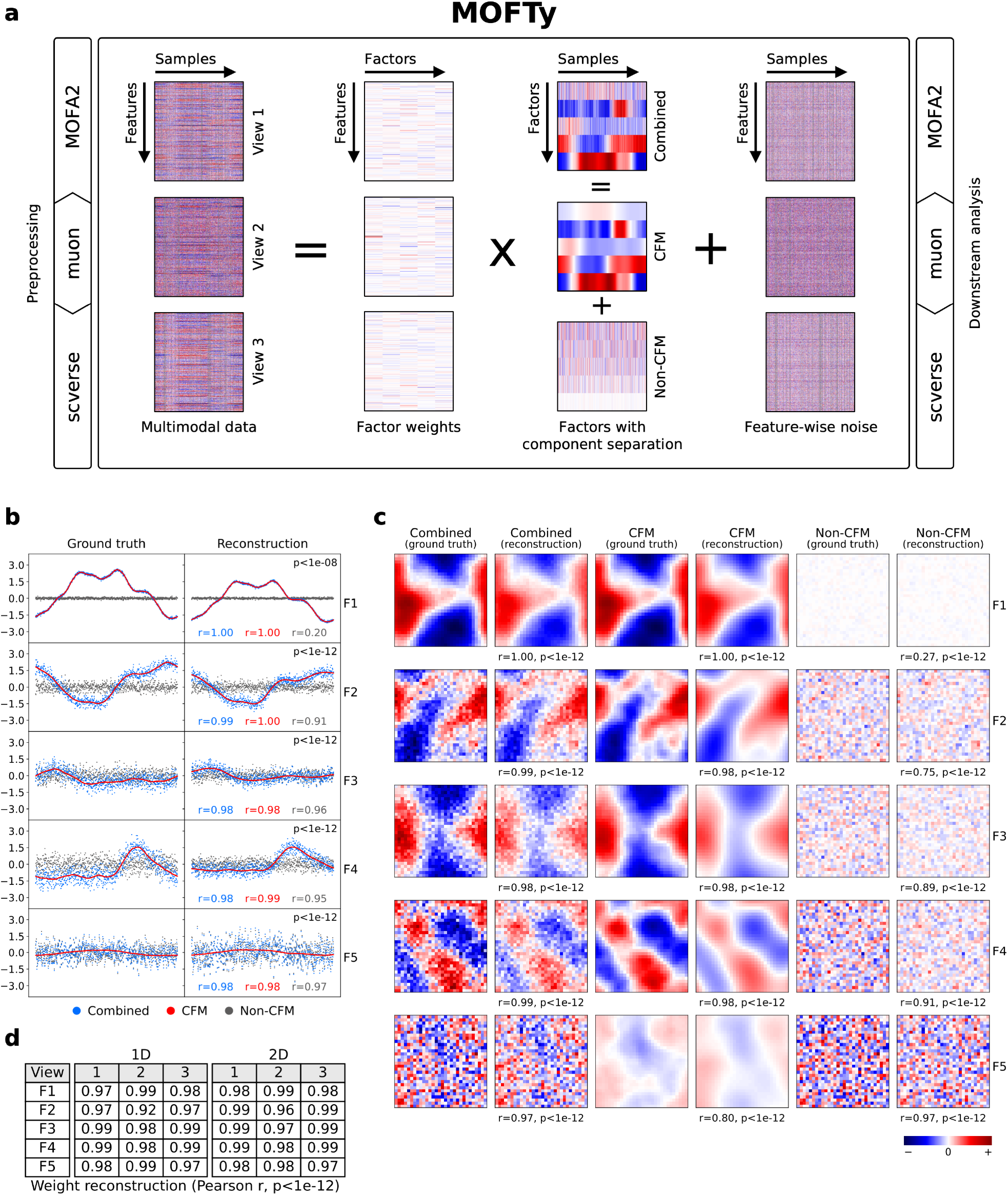
Overview and validation on synthetic data. **a**, Model overview illustrating the low-rank decomposition of a trimodal dataset along a one-dimensional (1D) continuous covariate using a Gaussian likelihood. Each of the three views is decomposed into the product of a view-specific weight matrix and a shared five-factor matrix, plus feature-wise noise. Each latent factor is further separated into a covariate-dependent correlated field model (CFM) component and a non-CFM component that captures variation not explained by the CFM. Gray indicates missing values in the data matrices. MOFTy integrates with MOFA2 and, via muon, with the scverse ecosystem for preprocessing (left subpanel) and downstream analyses (right subpanel), enabling interoperability with existing single-cell and multimodal analysis workflows. **b–d**, Validation on synthetic trimodal datasets with 1D and 2D covariates and known ground truth (five factors, F1–F5). Pearson correlation coefficients (*r*) with two-sided *p*-values between the ground truth and inferred weights/factors are reported. A comparison of ground truth and recovered factors is illustrated in **b** for a 1D example and in **c** for a 2D example. Diagrams in **b** and heatmaps in **c** display the factor values. For the 2D example in **c**, values of the combined, CFM, and non-CFM components are displayed at spatial coordinates using a shared normalized color scale across components. **d**, Inferred weights are compared to the ground truth weights for the 1D and 2D examples across all three views.

To facilitate preprocessing and downstream analyses, MOFTy integrates with MOFA2 [11–13, 30] and, through muon [31], with the scverse ecosystem [32], thereby enabling efficient data handling and interoperability with existing single-cell and multimodal analysis workflows. This is possible because MOFTy directly supports MOFA2- and muon-compatible data formats for both input and output.

We evaluated MOFTy in three stages to assess recovery of latent factor components, their interpretability, and practical utility across different data modalities. First, we used synthetic datasets with known latent structure to test whether MOFTy accurately reconstructs factors, weights, and the separation into CFM and non-CFM components. Second, we analyzed a publicly available spatial dual-omics dataset from human glioblastoma to illustrate how MOFTy resolves latent spatial structure into smooth spatial gradients and localized heterogeneity. Third, we applied MOFTy to a triple-omics mouse gastrulation dataset [33] that was previously analyzed with MEFISTO [13].

### 2.1 Validation on synthetic data

To illustrate the ability of MOFTy to recover latent factors with known CFM and non-CFM components (Fig. 1b–d, Ext. Data Fig. 4), we used the MOFTy forward model to generate five latent factors with varying contributions from CFM and non-CFM components in one dimension (900 points) and two dimensions (30*×*30 pixels). We next generated three view-specific weight matrices (300 features each), multiplied them by the latent factor matrix, added feature-specific Gaussian noise, and randomly masked 20% of the data columns in each view (Table 2, synthetic data generation; Methods, Section 4.8). To demonstrate the scalability of MOFTy, we also generated a larger 2D synthetic dataset with 100×100 pixels and 1,000 features per view (Methods, Section 4.8). We then trained MOFTy on these synthetic datasets using five latent factors, Gaussian likelihoods for all views, and the same model configuration as for all posterior approximations in this article (Methods, Section 4; Table 2, posterior approximation).

MOFTy recovered the factor weights and the combined factors accurately (Fig. 1b–d, Ext. Data Fig. 4). Recovery of the individual components tracked their share of the factor variance: components carrying a substantial share were recovered closely, whereas components with a very small share—such as the non-CFM component of factor 1—were recovered only weakly, as expected when a component contributes little to the factor it belongs to. The corresponding aligned posterior uncertainty estimates (Methods, Section 4.5; Supplementary Information) were consistent with the recovered factors and weights. Overall, uncertainty estimates were higher for the larger dataset with 100 × 100 pixels.

### 2.2 Decomposition of a spatial dual-omics dataset from human glioblastoma

Next, we applied MOFTy to a publicly available human glioblastoma dataset obtained from 10x Genomics, which was generated using the Visium CytAssist platform (Code and data availability, Section 5). By profiling whole-transcriptome gene expression together with targeted protein expression, this assay supports integrated characterization of tumor-associated and microenvironmental variation. After quality control and normalization, the dataset comprised 5,573 spatial spots, the top 2,000 highly variable genes, and 31 protein markers (Methods, Section 4.7.1). To guide the choice of the number of latent factors, we applied principal component analysis (PCA [34, 35]) to the standardized data matrices concatenated across features. Based on this analysis, in which the first four principal components each explained at least 2% of the variance and all subsequent components fell below this threshold, we initialized MOFTy with four latent factors using the model configuration described in Methods (Sections 4.2–4.7.1). After model training, the four resulting latent factors explained 23.7% and 45.9% of the variance in the gene and protein expression modalities, respectively. MOFTy also robustly recovered these four factors when initialized with a larger number of latent factors (Section 2.2.2).

The CFM components exhibited strong spatial autocorrelation (Moran’s *I* = 0.99), consistent with coherent spatial organization. In contrast, the non-CFM components showed weaker autocorrelation: factors 2–4 had positive Moran’s *I* values ranging from 0.20 to 0.41, indicating spatially localized, fine-scale variation not captured by the correlated field model, whereas factor 1 had approximately zero autocorrelation and a very low-magnitude non-CFM component (Fig. 2a). In both modalities, the CFM component accounted for the majority of explained variance, while the non-CFM component contributed substantially in factors 2–4 (Fig. 2b). Note that, unlike in PCA with orthogonality constraints, variance decomposition across factors is generally not additive in factor models, as factors need not be orthogonal. Variance decomposition across the CFM and non-CFM components is likewise non-additive: component-wise values are obtained by propagating each component through the shared loading matrix while holding the other at zero. The individual contributions can therefore be larger or smaller in sum than the variance explained by the combined factor, and a single component may reconstruct a given set of features worse than its feature-wise mean, yielding negative values.

**Fig. 2.**
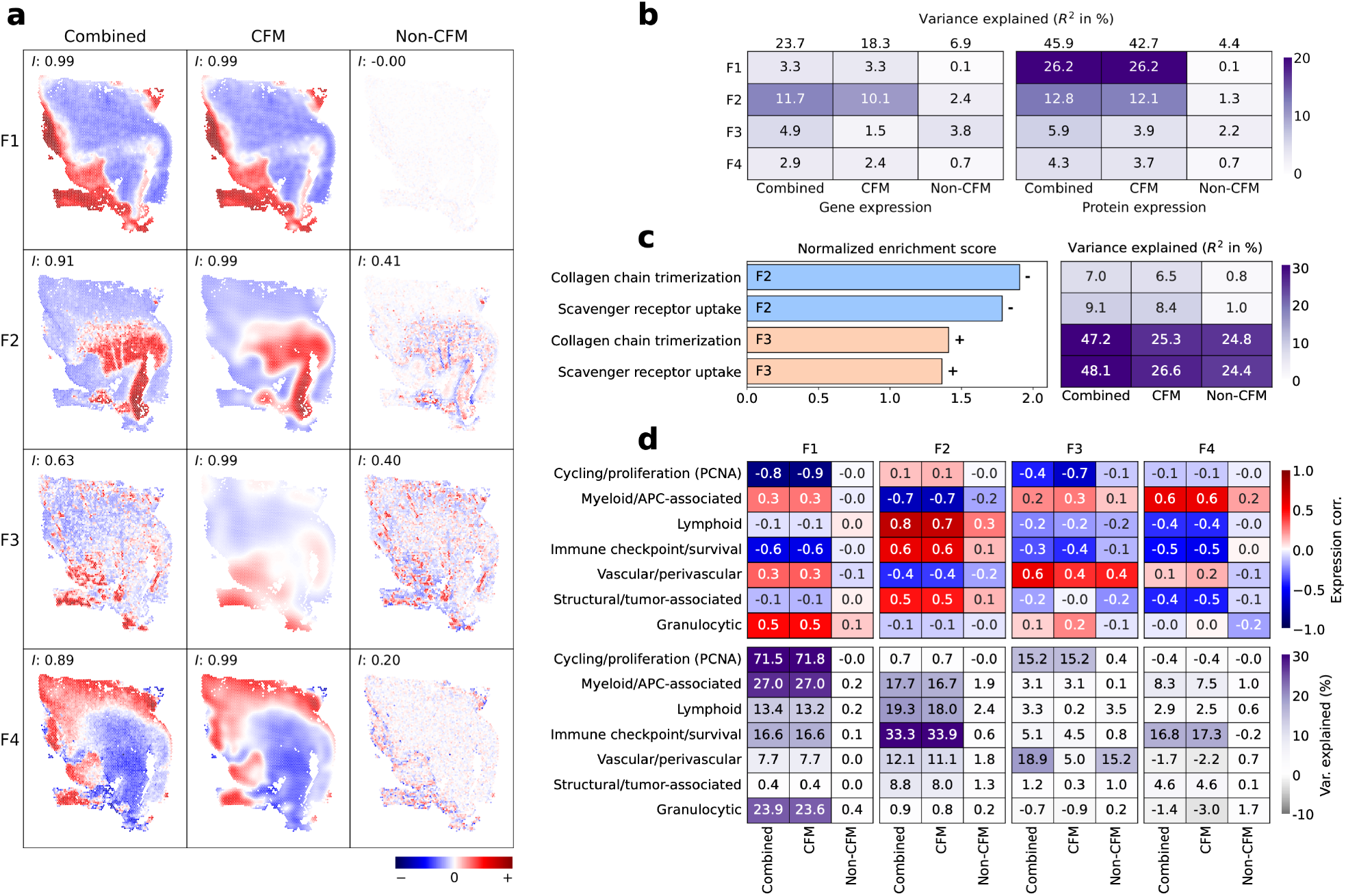
Decomposition of a spatial dual-omics dataset from human glioblastoma. **a**, MOFTy factor decomposition into CFM and non-CFM components (factors F1–F4). Spatial autocorrelation is quantified by Moran’s *I*, computed from the corresponding neighbor graph in scanpy [36]. For each factor, values of the combined, CFM, and non-CFM components are displayed over spatial spot coordinates using a shared normalized color scale across components. **b**, Variance explained (*R*^2^) decomposition across factors and modalities, where the variance explained by the combined, CFM, and non-CFM components is shown separately. Total variance explained by each factor is shown at the top of the corresponding column. **c**, Gene set enrichment analysis of gene expression modality weights across factors, together with the variance explained by the specified factor component in each corresponding gene set. The sign indicates the direction of association with the factor. **d**, Protein marker set association profiles and explained (*R*^2^) variance across factors. Negative variance explained values indicate that the corresponding factor component reconstructs the marker set worse than its feature-wise mean. The color scales represent variance explained in % and the correlation (Pearson *r*) between mean expression profiles within each marker set and factor components (two-sided *p*-value *<* 10^−12^ for *r* > 0.1). Correlations were classified as follows: 0.2 ≤ |*r*| *<* 0.4 (weak), 0.4 ≤ |*r*| *<* 0.6 (moderate), 0.6 ≤ |*r*| *<* 0.8 (strong), |*r*| ≥ 0.8 (very strong).

To interpret the inferred factors, we performed pathway enrichment analyses of the factor weights in the gene expression modality using GSEApy [37] and Reactome [38] (Fig. 2c, Ext. Data Fig. 6–7). Furthermore, we assessed the top features for each factor (by absolute weight) together with the sign of their weights (Ext. Data Fig. 5). At the protein level, we examined protein-factor association profiles (Fig. 2d) via predefined marker sets representing canonical cell populations and functional components of the tumor microenvironment [39–44]. For consistency with the assay output, the following marker set definitions use the feature names reported by the 10x Genomics protein panel:

1. Cell cycle and proliferation: PCNA.
2. Myeloid and antigen-presenting cell (APC)-associated: CD163, CD68, CD14, ITGAM, ITGAX, FCGR3A, HLA-DRA, PTPRC-1 (CD45RA), and PTPRC-2 (CD45RO).
3. Lymphoid: T-cell markers (CD3E, CD4, CD8A, CD27, CCR7), B-cell-associated markers (MS4A1, CD19, PAX5, CR2), and additional adaptive immune markers (CXCR5, CD40, PDCD1).
4. Immune checkpoint and survival-associated: CD274 (PD-L1) and BCL2.
5. Vascular and perivascular: PECAM1, ACTA2, and VIM.
6. Structural and tumor-associated: EPCAM, KRT5, and SDC1.
7. Granulocytic: CEACAM8.

In the gene expression modality, factor 1 highlighted the hemoglobin genes *HBA2* and *HBB* among the top feature weights (both positive). Elevated hemoglobin transcripts can indicate erythrocyte- or blood-associated signal, suggesting blood-rich, intravascular, or hemorrhagic tissue regions. Gene set enrichment analysis identified pathways with nominal enrichment, but the evidence was weak after multiple-testing correction, with all FDR-adjusted *q*-values above 0.6. At the protein level, factor 1 showed a very strong negative correlation with PCNA, a well-established marker of DNA replication and cellular proliferation [39] that has also been used to assess proliferative activity in brain tumors, including gliomas [45]. Marker set analysis further revealed a moderate positive correlation with the granulocytic marker CEACAM8 and strong negative correlations with immune checkpoint/survival-associated markers. Across protein marker sets, variance explained by factor 1 was dominated by the CFM component (Fig. 2b). Together, these findings indicate that factor 1 captures a spatially continuous axis associated with blood- or erythrocyte-related signal and granulocytic marker expression, inversely related to proliferative activity.

Factors 2 and 3 captured distinct spatial organizations but shared overlapping transcriptional features, pathway enrichments, and protein-marker correlations. However, these shared associations were embedded in different factor-specific molecular contexts. Gene weights for factor 2 were dominated by neuronal and synaptic genes (all positive), including *VSNL1, SNAP25, CREG2, OLFM1*, and *ATRNL1*. Gene set enrichment analysis supported this pattern: pathways enriched at the positive end of factor 2 were related to neuronal signaling, sensory processing, neurotransmission, GABA receptor activation, and metabolic or anti-inflammatory signaling, whereas pathways enriched at the negative end were dominated by extracellular matrix (ECM) and uptake-related processes (Ext. Data Fig. 6). By contrast, factor 3 highlighted collagen and ECM-related genes among the top factor weights (all positive), including *COL3A1, COL1A2, COL1A1, DCN*, and *FN1*, with positively enriched pathways centered on collagen/ECM remodeling and uptake-related activity. The different signs of gene weights and pathway enrichments should not be interpreted as a direct biological reversal between factors, because signs are defined relative to the orientation of each individual factor.

We highlighted two pathways enriched in both factors: collagen chain trimerization, and binding and uptake of ligands by scavenger receptors (Fig. 2c, Ext. Data Fig. 6). In factor 2, both pathways were enriched at the negative end and opposed positive neuronal and synaptic gene weights, positioning this factor as a contrast between ECM-rich and neuronal tissue compartments. In factor 3, both pathways were enriched at the positive end and aligned with positive collagen and ECM gene weights, without an opposing neuronal signature. The two factors therefore capture the same ECM- and uptake-associated processes in different configurations: as one pole of a tissue-compartment axis in factor 2, and as a self-contained ECM signature in factor 3. This distinction is also reflected in the variance decomposition: factor 2 explained less variance in the highlighted pathways than factor 3, and its contribution was predominantly attributable to the CFM component, whereas in factor 3 both components contributed substantially, indicating that the ECM signature carries localized structure beyond a smooth spatial gradient.

At the protein level, factor 2 showed strong positive correlations with lymphoid and immune checkpoint/survival-associated marker sets, with variance explained predominantly by the CFM component (Fig. 2d). By contrast, myeloid/APC-associated marker sets showed a strong negative correlation with factor 2. Vascular/perivascular markers also showed a moderate negative correlation with factor 2, with variance explained predominantly by the CFM component. Thus, factor 2 defined a predominantly spatially continuous axis linking neuronal-like transcriptional features with lymphoid and immune-checkpoint-associated protein signals, while opposing regions were characterized by myeloid/APC-associated, vascular/perivascular, and ECM-associated features. Factor 3, in contrast, showed a strong positive correlation with vascular/perivascular marker sets and a strong negative correlation with PCNA (only in the CFM component). Unlike factor 2, where variance explained for vascular/perivascular markers was dominated by the CFM component, in factor 3 these markers were largely attributable to the non-CFM component, which retained positive spatial autocorrelation, consistent with spatially localized structure within vascular/perivascular niches. Together, factors 2 and 3 describe related processes in different spatial regimes: a predominantly continuous axis in factor 2, and a more heterogeneous niche pattern in factor 3.

Gene set enrichment analysis in factor 4 (Ext. Data Fig. 7) yielded a broad pathway profile without a clear theme, so we focused the interpretation on leading gene weights and protein marker associations. The top gene weights in factor 4 suggested a mixed glial-myeloid program, including lipid- and complement-related genes as well as myelin- and microglial/macrophage-associated markers (*APOC1, PLTP, C1QA, PLP1, GPNMB, MS4A7*; all positive). This pattern is consistent with a glial- and immune-microenvironment-associated transcriptional profile rather than a purely oligodendrocyte-lineage signature. At the protein level, factor 4 explained a substantial fraction of variance in the immune checkpoint/survival-associated marker set, with a moderate negative correlation, and a modest fraction of variance in the myeloid/APC-associated marker set, with a strong positive correlation. Together, these results suggest that factor 4 captured a mixed glial-myeloid microenvironmental signature, linking myelin-related glial features with macrophage/microglial lipid- and complement-associated features.

To illustrate how latent factor-based feature reconstruction can be used to denoise expression patterns and disentangle spatial organization from local heterogeneity, we reconstructed the features with the highest absolute weights for factors 1–4 in both modalities (Ext. Data Fig. 8). Overall, MOFTy robustly reconstructed the dominant expression patterns. The patterns reconstructed from the combined component were less noisy than the observed data, consistent with denoising arising from the low-rank representation and modeling of feature-specific noise levels. Reconstructions based on the CFM component captured gradual, smoothly varying spatial patterns, whereas those from the non-CFM component retained fine-scale variation and localized heterogeneity.

#### 2.2.1 Comparison with PCA and MEFISTO

To contextualize the spatial structures inferred by MOFTy, we compared the four MOFTy factors (Fig. 2a) with four factors inferred by MEFISTO [13] and with the scores of the first four principal components obtained by applying PCA to the concatenated preprocessed modalities (Ext. Data Fig. 9, Ext. Data Fig. 10). PCA scores showed similarities to the MEFISTO and MOFTy factors but appeared noisier than both the MEFISTO factors and the combined MOFTy representation.

Unlike MOFTy and MEFISTO, PCA does not provide GP-based inference or an explicit noise model; instead, it constrains components to be orthogonal while maximizing total variance. Consequently, directions with large noise variance can enter the leading components alongside true latent structure. This can cause leading PCA components to mix smooth spatial structure with non-smooth or noisy variation and can split biologically related signals across orthogonal directions.

For MEFISTO, we evaluated configurations with 1,000 and 3,000 inducing points, as well as a full GP configuration using all 5,573 spatial spots. As expected, the full GP configuration showed the greatest similarity to the combined MOFTy factors, as reflected by very strong correlations (*r* ≥ 0.98) between the corresponding factors and weights (Ext. Data Fig. 9a). Across factors, the full GP configuration tended to explain slightly more variance than the combined MOFTy factors in the gene expression modality, whereas the opposite pattern was observed in the protein expression modality (Ext. Data Fig. 9b,c).

At this point, we emphasize that a key contribution of MOFTy is not to produce a different factorization, but rather to decompose each latent factor into a CFM component and a complementary non-CFM component.

Comparison of the MEFISTO factors across the three different configurations revealed two main effects. First, the observed spatial smoothness decreased as the number of inducing points increased from 1,000 to 3,000, and decreased further under the full GP configuration, as indicated by lower Moran’s *I* values across factors. Second, the factor-wise inferred GP variance-scale parameter increased with the number of inducing points. These apparently opposing trends reflect the distinction between the learned variance fraction and the spatial autocorrelation of the resulting factor realizations. In sparse inducing-point approximations, the posterior is represented through inducing variables at a finite set of locations, while factor values elsewhere are obtained through kernel-based GP conditioning. A more limited inducing representation can therefore suppress local spatial variation between inducing locations, while the free variational parameters at the inducing points themselves pull the posterior toward the local data, yielding scattered point-like features on an otherwise over-smoothed field. In datasets such as the present one, where the latent factors exhibit substantial spatial heterogeneity, a richer GP representation allows non-smooth, fine-scale variation to be expressed throughout the factor realizations rather than only at a finite set of inducing locations.

To compare the factor decomposition of MOFTy with a parametric kernel approach, we applied the interpolation functionality provided in MOFA2 [30] to the full-GP MEFISTO factors to obtain the smooth GP component at the observed coordinates (Ext. Data Fig. 11a). We used this interpolation to construct an additive decomposition analogous to the one in MOFTy: the interpolated component plays the role of the CFM component, and the difference between the inferred and interpolated factors plays the role of the non-CFM component. However, the decompositions are not equivalent. We observed that the difference between the original and interpolated factors showed weak negative spatial autocorrelation across all factors (Moran’s *I* = −0.16 to −0.05). Alternating-sign patterns are characteristic of smoothing residuals rather than coherent spatial structure. This observed pattern was also consistent with the short length scales inferred by MEFISTO for the GP kernel (Ext. Data Fig. 11a).

In MOFTy, the corresponding fine-scale variation was instead captured by a positively autocorrelated non-CFM component in factors 2–4 (Moran’s *I* = 0.20 to 0.41). In factor 1, the CFM component accounted for essentially all spatial structure, while the non-CFM component was negligible and spatially uncorrelated (Moran’s *I* ≈ 0).

The variance decomposition showed the same pattern (Ext. Data Fig. 11b). For factors 2–4, the non-CFM component explained more variance than the difference between the original and interpolated MEFISTO factors, with the largest gap observed for factors whose non-CFM components showed the strongest spatial structure.

We also compared the performance and complexity of MEFISTO and MOFTy, both trained on a GPU (Methods, Section 4.6; Table 4). Per model, MOFTy required substantially less combined CPU and GPU memory than MEFISTO with a full GP. Since MOFTy models were trained concurrently in sets of four, per-model values provide a direct comparison. Per model, MOFTy also required less combined memory than MEFISTO with 1,000 inducing points. The runtime of the full-GP MEFISTO configuration was 20% shorter than that of MOFTy, which likely reflects, at least in part, our concurrent training of four MOFTy models from different random initializations and the use of geoVI [28]. Unlike the factorized variational posterior used for non-GP parameters in MEFISTO, geoVI preserves dependencies and captures local posterior curvature (Methods, Section 4.4).

We note that all MOFTy downstream analyses were based exclusively on CPU-trained models, demonstrating that MOFTy remains practical without GPU access—owing to the memory efficiency and optimized CPU execution of NIFTy—although CPU training took roughly twice as long as GPU training (Methods, Section 4.6; Table 4).

#### 2.2.2 Robustness of latent factors and correlation structures

We assessed the robustness of the inferred latent representation by computing aligned posterior uncertainty estimates, which indicated that the inferred structures were stable across seeds, with factor weights well constrained by the data (Methods, Section 4.5; Supplementary Information). For both the CFM and non-CFM components, uncertainty estimates tended to be higher when their contributions to the combined factor were smaller.

For each MOFTy factor, we computed the diagonal approximation of the aligned posterior power spectra of the factor-wise CFM components to characterize the inferred latent correlation structures (Ext. Data Fig. 14a; Methods, Section 4.5). Across factors, the power spectra showed a clear decay from low to high frequencies, consistent with the strong spatial autocorrelation of the CFM components (Moran’s *I*; Fig. 2a). The inferred spectra exhibited shapes that cannot be captured by single-length-scale parametric kernels, such as the stationary and isotropic RBF and Matérn kernels used in existing approaches [13–15, 46]. Factors 1, 2, and 4 exhibited multi-scale structure with shoulders and multiple changes in slope, whereas factor 3 followed an approximate power law across the full frequency range. All four factors additionally showed small-scale irregularities.

We further assessed the robustness of the inferred latent factors across different configurations with six, eight, and ten factors (Ext. Data Fig. 12). Latent factors and component separation were consistently recovered across configurations. The CFM components showed very strong correlations (|*r*| > 0.90), with one exception (|*r*| = 0.84). For the non-CFM components, correlations were generally strong (|*r*| ≥ 0.70), with two exceptions (|*r*| = 0.66 and |*r*| = 0.37) in factors with low contributions from the non-CFM component. The non-CFM components of factors 2 and 3 in the four-factor configuration were consistently recovered across all configurations with |*r*| ≥ 0.95. For the factor weights, we observed very strong correlations in the gene expression modality (|*r*| ≥ 0.90), with four exceptions (0.80 ≤ |*r*| *<* 0.90), and very strong correlations in the protein expression modality (|*r*| ≥ 0.80), with three exceptions (0.69 ≤ |*r*| *<* 0.80).

### 2.3 Decomposition of a triple-omics mouse gastrulation dataset

Finally, we applied MOFTy to a single-cell mouse gastrulation scNMT-seq dataset [33], which was previously analyzed with MEFISTO [13], providing a useful reference for assessing whether MOFTy recovers similar patterns. The dataset was obtained from the corresponding MEFISTO tutorial [30] and comprises 1,518 cells profiled across three omic layers: RNA expression (887 features), DNA methylation (500 features), and chromatin accessibility (500 features). The two epigenetic modalities are sparse, with methylation and accessibility measurements available for only 33.8% and 29.9% of the cells, respectively. We note that the MEFISTO publication reports 945 features for the RNA modality, whereas the tutorial provides only 887 features. This discrepancy may reflect differences in the underlying data or in the software versions used for preprocessing.

We used the same two-dimensional UMAP [47, 48] coordinates provided in the MEFISTO tutorial, treating them as a two-dimensional covariate for the CFM components of the MOFTy factors (Methods, Section 4.7.2). Each point represents a single cell, colored according to the lineage annotations from the original study [33] (Fig. 3a). We initialized MOFTy with ten factors, consistent with the MEFISTO tutorial, and retained all factors after training rather than restricting the analysis to factors explaining at least 1% of the variance in the RNA modality.

**Fig. 3.**
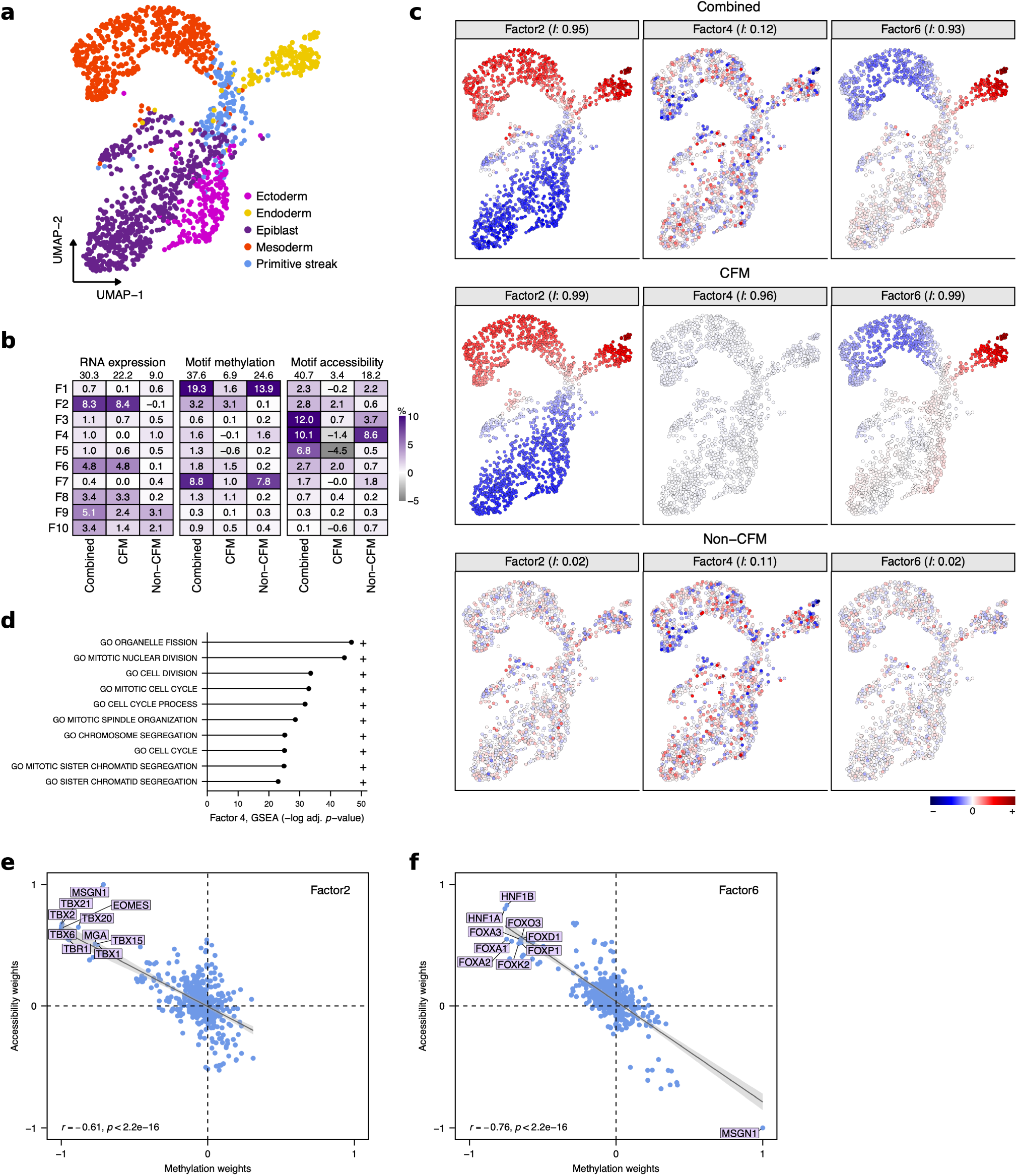
Decomposition of a triple-omics mouse gastrulation dataset. **a**, UMAP [47, 48] embedding derived from the RNA modality, colored by lineage assignments [33], as used in the MEFISTO publication [13] and the corresponding tutorial [30]. **b**, Contributions to variance explained (in %) across factors (F1–F10) and modalities (RNA, DNA methylation, and chromatin accessibility), decomposed into combined, CFM, and non-CFM components. Negative variance-explained values indicate that the corresponding factor component reconstructs the modality worse than its feature-wise mean. **c**, Factor decomposition into CFM and non-CFM components. Factor values for F2, F4, and F6 across the combined, CFM, and non-CFM components are displayed over UMAP coordinates using a shared normalized color scale for each factor. Spatial autocorrelation is quantified by Moran’s *I*, computed from the UMAP coordinates using a 53 *×* 65 queen-contiguity grid corresponding to the unpadded grid size of the correlated field configuration (Methods, Section 4.7.2). **d**, Gene set enrichment analysis of RNA modality weights for factor 4, showing the top ten enriched pathways from the MSigDB database [49], ranked by −log_10_ of the FDR-adjusted *p*-value. **e, f**, DNA methylation vs. chromatin accessibility weights for factors 2 and 6. Highlighted are the top 10 features ranked by the sum of their absolute weights across the two modalities. Solid lines show least-squares fits with 95% confidence intervals (gray). Pearson correlations (*r*) with two-sided *p*-values are reported. The sign indicates the direction of association with the factor.

Overall, the MOFTy factors explained 30.3%, 37.6%, and 40.7% of the variance in the RNA, DNA methylation, and chromatin accessibility modalities, respectively (Fig. 3b). In the RNA modality, most variance was explained by the CFM component, whereas in DNA methylation and chromatin accessibility it was explained predominantly by the non-CFM component.

In factor 4, the explained variance was mainly attributable to the non-CFM component, with RNA modality weights enriched for cell-cycle-associated gene sets (Fig. 3d; MSigDB [49]). Factors 2 and 6 exhibited the expected anti-correlation between DNA methylation and chromatin accessibility factor weights (Fig. 3e,f; [50]). Among the top factor weights, MOFTy identified transcription factors associated with mesoderm fate, including TBX6 and MSGN1, in factor 2 (Fig. 3e), and transcription factors associated with endoderm fate, including FOXA2 and HNF1A/HNF1B, in factor 6 (Fig. 3f). The patterns observed for factors 2, 4, and 6 were consistent with those previously reported using MEFISTO [13, 30].

As in the MEFISTO publication, we reconstructed feature values for the transcription factors MSGN1 and FOXA2 in the DNA methylation and chromatin accessibility modalities to illustrate denoising and missing-value imputation (Ext. Data Fig. 13a,b). Reconstructions based on the CFM component captured gradual, smoothly varying spatial patterns, whereas those from the non-CFM component retained fine-scale variation. The observed high spatial autocorrelation (Moran’s *I* = 0.97 to 0.99) of CFM-based reconstructions was qualitatively consistent with the interpolation results reported in the MEFISTO publication, which were obtained by interpolating factor values at the observed coordinates using the RBF kernel with the learned factor-specific length scales and variance fractions.

Aligned posterior uncertainty estimates for factors and weights indicated that the latent structures were stable (Methods, Section 4.5; Supplementary Information). The high sparsity of the DNA methylation and chromatin accessibility modalities was reflected in higher factor weight uncertainty than for RNA. The diagonal approximation of the aligned power spectra showed higher uncertainty than in the glioblastoma dataset (Ext. Data Fig. 14b; Methods, Section 4.5). This is consistent with the weaker constraint on the power spectra arising from the irregular distribution of the UMAP-derived coordinates and the extensive missing data in the DNA methylation and chromatin accessibility modalities. Across factors, the inferred spectra exhibited diverse shapes, with multiple changes in slope and small-scale irregularities.

## 3 Discussion

We present MOFTy, a Bayesian Gaussian process (GP) factor analysis framework for integrating multimodal data with continuous covariates. MOFTy replaces pre-specified GP kernel families with the correlated field model (CFM) implemented in NIFTy [20–25], in which the covariance structure of each latent factor is inferred through a nonparametric power spectrum. Each factor is further separated into a CFM component, which captures the correlated structure described by this spectrum, and a complementary non-CFM component. To our knowledge, MOFTy is the first Bayesian multimodal GP factor analysis framework to combine such an explicit factor-level separation with a nonparametric power spectrum model. In the glioblastoma dataset, the inferred power spectra exhibited multi-scale structure with shoulders and multiple changes in slope (Ext. Data Fig. 14a), shapes that the single-length-scale RBF and Matérn kernels used in existing frameworks [13–15, 46] cannot represent, indicating that the nonparametric spectral model resolves correlation structure inaccessible to fixed kernel families.

We found that component separation is particularly informative in settings where structured biological gradients coexist with localized heterogeneity. In a spatial dual-omics dataset from human glioblastoma with paired gene and protein expression measurements, MOFTy resolved tumor-associated, immune-microenvironmental, extracellular matrix, and signaling axes across both modalities. Broad, spatially coherent patterns were captured primarily by the CFM component, whereas the non-CFM component contributed more strongly to spatially localized variation in specific contexts, most notably extracellular matrix remodeling and vascular or perivascular niches. This separation reflects a distinction between large-scale tissue organization, which varies along extended spatial gradients, and microenvironmental niche structure, which is confined to more localized regions. When MEFISTO [13] was trained on this glioblastoma dataset, the learned length scales of the factor-wise GP kernels were only a few times the spot spacing, and the difference between the factors and their smooth interpolation was weakly anti-correlated, indicating that tissue-scale and fine-scale variation were absorbed into a single smooth field.

In a triple-omics mouse gastrulation dataset with extensive missing data, MOFTy recovered cross-modal patterns consistent with those reported for MEFISTO [13], indicating that the added flexibility does not come at the cost of recovering known structure.

MOFTy has several limitations. First, the CFM component assumes a homogeneous and isotropic correlation structure a priori, so spatially varying or directional correlation patterns are not accommodated by this prior. Leveraging the flexibility of the NIFTy framework, the model could be extended and generalized to inhomogeneous and anisotropic correlation structures, for example by interleaving and combining multiple correlated field components into a unified operator. Second, although non-negativity constraints can be imposed on weights and combined factors, component separation is not yet available when these constraints are applied. Third, MOFTy does not provide native support for multi-group settings [12–14], guided factors [14, 51], or the incorporation of prior domain knowledge [14, 52].

State-of-the-art single-cell and spatial omics technologies are generating datasets at rapidly increasing scales, from hundreds of thousands to millions of cells or spatial locations. While MOFTy already harnesses the scalable correlated field model in NIFTy, further computational advances will be needed to extend GP factor analysis to spatial omics datasets beyond megapixel resolution. NIFTy offers a promising route toward this goal through iterative charted refinements (ICR [53]), a hierarchical multi-resolution refinement scheme that avoids storing the full covariance. ICR has been successfully applied to three-dimensional GPs with up to 122 billion degrees of freedom [54] and outperforms state-of-the-art methods like KISS-GP [55] by an order of magnitude in computational speed [53]. This line of work has recently been extended by GraphGP [56], which uses a graph-based Vecchia approximation to support arbitrary point distributions while retaining the scalability of ICR. We envision MOFTy as a new framework for scaling GP factor analysis to new and increasingly large spatial omics datasets, transforming massive-scale molecular profiles into interpretable maps of spatial biological organization.

## 4 Methods

The Methods section is organized into eight subsections. We first provide an overview of related methods in Gaussian process (GP) factor analysis (Section 4.1). We then describe the MOFTy model design (Section 4.2) and the construction of the correlated field model (Section 4.3). Next, we summarize the variational inference scheme, including the optimization and initialization strategy (Section 4.4). Section 4.5 describes the alignment of posterior samples across random initializations. We then discuss the computational complexity and scalability of MOFTy and the NIFTy framework (Section 4.6). Finally, we describe the preprocessing steps for the glioblastoma and mouse gastrulation datasets (Section 4.7), along with the synthetic data generation procedure (Section 4.8).

### 4.1 Related methods in Gaussian process factor analysis

In this subsection, we contrast three lines of related work that are relevant for positioning MOFTy: multi-modal Gaussian process factor analysis (GPFA), frequency-domain GP approximations, and GPFA models that separate GP and non-GP factors and/or support non-negativity constraints.

Gaussian processes are a key concept in modern machine learning for nonparametric regression modeling and function estimation [57]. They have been integrated into factor analysis frameworks [13–15, 46, 58–60] to capture latent structures of high-dimensional (biological) datasets along continuous covariates. Across these frameworks, the covariate-dependent GP components are typically specified using parametric, stationary, isotropic kernels; common choices are the radial basis function (RBF) and Matérn kernels. For each factor, the kernel is governed by a learnable amplitude and a single length scale and, in the case of Matérn kernels, by an additional smoothness parameter fixed before training. By construction, kernels of this form cannot represent correlations characterized by multiple distinct scales, spectra with multiple independently varying slope regimes, or localized spectral irregularities—structures that can be represented by the nonparametric power-spectrum model in NIFTy [20– 25]. In frameworks such as MEFISTO [13], which supports RBF kernels, and, more recently, MOFA-FLEX [14], which additionally supports Matérn kernels, the selected kernel is augmented with an uncorrelated factor-specific term. A factor-specific variance-scaling parameter determines the relative contributions of the correlated and uncorrelated components. Because the uncorrelated term contributes frequency-independent power, it does not change the characteristic shape of the combined spectrum.^1^

In practice, the flexibility of the inferred GPs is often further restricted by sparse variational methods that rely on inducing-point or low-rank approximations of the covariance [18, 19]. For large datasets, sparse approximations are often unavoidable, as the covariance matrix grows quadratically in memory with the number of grid points, while operations such as inversion or Cholesky factorization scale cubically. Training MEFISTO on the glioblastoma dataset with sparse configurations produced fields with isolated, point-like features against an over-smoothed background (Ext. Data Fig. 10). This is consistent with reconstruction from a sparse set of locations: the field remains close to the data at the inducing points but is over-smoothed between them when the inducing-point density is low relative to the data resolution.

Even with a full GP, a single learned length scale per factor can force a compromise between large-scale and fine-scale variation. At the level of field realizations, this became visible in the glioblastoma dataset when we applied the interpolation functionality of MEFISTO [13, 30]: evaluated at the observed covariates, this interpolation returns the posterior mean of the smooth GP component given the factor values, equivalent to a factor-wise linear smoothing operator whose cutoff is set by the learned length scale and variance fraction. When that length scale is only a few times the spot spacing, as we observed in the glioblastoma dataset (Ext. Data Fig. 11a), this cutoff lies near the sampling resolution. Almost all resolvable variation was therefore retained in the smooth component, and the difference between the factor and its smooth component contained only high-frequency, anti-correlated variation.

This is in contrast to MOFTy, where the correlated field model (CFM) components and non-CFM components are modeled as separate latent fields. For the CFM component, the underlying correlated field model in NIFTy assumes the correlation structure to be homogeneous a priori, so by the Wiener–Khinchin theorem [26, 27] the two-point correlation function is diagonal in harmonic space; the additional isotropy assumption implies that the power spectrum depends only on the magnitude of the harmonic mode and not on its direction. The resulting one-dimensional function is assigned a flexible power-spectrum prior, consisting of a power-law trend with deviations modeled by an integrated Wiener process. Hyperpriors control the amplitude, average slope, flexibility, and roughness of the power spectrum (Section 4.3). The slope prior is typically chosen to favor decaying spectra, resulting in more regular field realizations. Neglecting uncertainty in the power spectrum—for instance by using a point estimate—introduces a “perception threshold” in which modes with low inferred variance, typically at high frequency, are damped toward zero even when they carry signal [61]. The correlated field model treats the field and its power spectrum jointly as random fields, so that the correlation structure can be inferred with uncertainty. In the configuration used here, MOFTy point-estimates the scalar spectral hyperparameters for numerical stability. Posterior samples are retained for the integrated Wiener process increments, which carry the deviations of the log-log power spectrum from its power-law trend and therefore the nonparametric shape of the spectrum. In frameworks such as MEFISTO and MOFA-FLEX, the correlation structure is instead fully specified by a small number of kernel parameters, all of which are point-estimated.

Frequency-based representations in harmonic space have recently been combined with inducing variables to improve the scalability of Gaussian-process approximations [62]. Such approaches have been applied to GPFA, including frequency-domain approximations [63] and non-conjugate GPFA with Poisson observations (SNP-GPFA [64]). These methods exploit the diagonalization of GP covariances in harmonic space, but treat the spectral representation primarily as a computational tool, with the underlying GP kernel family chosen in advance.

The unimodal non-negative spatial factorization hybrid model NSFH [15] separates spatial (GP) and non-spatial (non-GP) factors by summing two matrix factorizations, each with its own feature weights. SNP-GPFA [64], developed for time-series analysis of neural population activity, uses a similar construction with distinct loading matrices. Assigning distinct loadings identifies the separation from the data, at the cost of a separate factor count for each component and the loss of a direct correspondence between the components of a given factor. In MOFTy, the CFM and non-CFM components share a loading matrix, so the observation model constrains only their sum. This shared loading is what allows the two components to be interpreted as contributions to a single latent factor. While the relative variance of the two components and the inferred correlation structure are identified by the likelihood, the assignment of individual observations to the two components retains irreducible posterior uncertainty that MOFTy quantifies but cannot reduce with additional data.

Another aspect is compatibility with non-negativity constraints, which are supported by FISHFactor [46], NSFH [15], and MOFA-FLEX, but not by MEFISTO. MOFTy supports non-negativity constraints, with or without using a CFM prior for the factors. However, when non-negativity constraints are used together with a CFM prior, the constraints are applied to the combined latent factors, rather than separately to the non-CFM and CFM components. Although imposing the constraints separately is straightforward from an implementation perspective, the methodological justification for doing so is unclear, and whether a more principled construction would be preferable remains an open question for future work.

### 4.2 Model design

MOFTy integrates multiple observation models within a unified multimodal framework. At a high level, the model comprises three layers: an observation model that links each view to a shared low-rank predictor; heavy-tailed shrinkage priors on the view-specific factor weights; and a prior on the latent factors that, in the covariate-aware setting, decomposes each factor into a correlated field model (CFM) component and a complementary non-CFM component. We describe each layer below.

Let *Y* ^(1)^, …, *Y* ^(*M*)^ be data matrices across *M* views, with 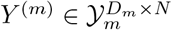, where *D*_*m*_ denotes the number of features in view *m* and *N* denotes the number of samples shared across all views. Here, *Y*_*m*_ denotes the observation space of view *m*, for example ℝ for Gaussian real-valued observations, *{*0, 1} for Bernoulli binary observations, and non-negative integers ℤ_≥0_ for Poisson count observations. Given *L* latent factors, MOFTy represents each view through a low-rank linear predictor

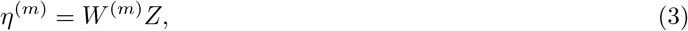

where 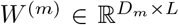 is the view-specific feature loading matrix and *Z* ∈ ℝ^*L*×*N*^ is the latent factor matrix shared across views. The linear predictor *η*^(*m*)^ is mapped to the parameters of the view-specific likelihood through an appropriate link function.

We first consider the standard factor analysis setting,

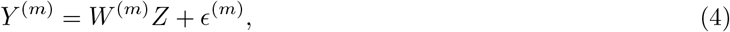

where 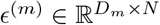 denotes independent additive Gaussian noise with mean zero. As in existing multi-omics factor analysis frameworks [11–14], we furthermore assume heteroskedasticity across features, i.e.,

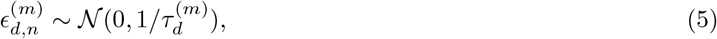

where 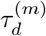 denotes the precision (inverse variance) for feature *d* in view *m*. This leads to the following Gaussian likelihood:

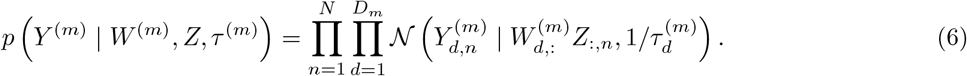

For binary observations, MOFTy uses a Bernoulli likelihood

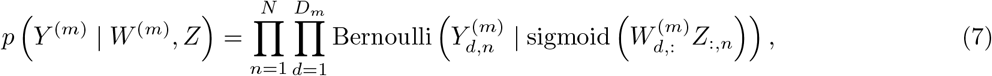

where the sigmoid (logistic link) function maps the linear predictor to the interval (0, 1). This Bernoulli formulation is generally incompatible with simultaneous non-negativity constraints on both weights and factors, because such constraints restrict the Bernoulli probabilities to the interval [0.5, 1). For count observations, MOFTy uses a Poisson likelihood

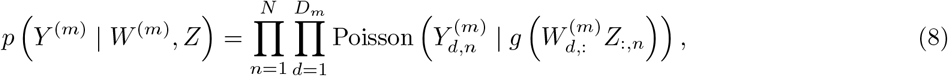

where *g* denotes an exponential or softplus [65] link function that maps the linear predictor to a positive rate parameter. Alternatively, when non-negativity is enforced on factors and weights (e.g., via the softplus or exponential transformations supported in MOFTy), an identity link can be used.

Likelihoods across different views are assumed to be independent, so the overall likelihood factorizes into a product across views. Like other probabilistic models, MOFTy naturally handles missing values without prior imputation: because the likelihood factorizes over observations, unobserved entries are simply omitted from the product. Posterior predictions for missing entries are obtained from the trained model itself, conditional on the observed entries and the priors on *W* ^(*m*)^ and *Z*.

Next, we specify the prior structures for the weights and factors. In NIFTy [20–25], all models are constructed within a generative, operator-based framework that starts from standardized variables *ξ* ∈ ℝ^*A*^, with *ξ* ~ *N* (0, I_*A*_). Here, *A* = (*a*_1_, …, *a*_*n*_) defines the shape of a vector (*n* = 1) or the shape of an array (*n* > 1), and I_*A*_ denotes the identity covariance matrix. The model variables are obtained from *ξ* through a deterministic forward model; posterior samples are propagated by applying this map to posterior samples of *ξ*. For further details, we refer to the original MGVI and geoVI publications [28, 29], which provide a comprehensive introduction to these concepts and their implementations in NIFTy.

The goal is to construct a forward model that encodes our prior assumptions about the latent factors, their correlated field structure, and the factor weights. Given a probability density *p*, we denote by *p*^*o*^ the operator that transforms standard normal random variables into random variables distributed according to *p* via cumulative distribution function (CDF) transformations, i.e.,

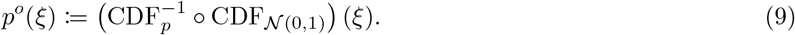

For arrays of standardized variables, the operator *p*^*o*^ is defined element-wise. If the inverse CDF of *p* is not available in closed form, we use numerical approximations to construct the operator.

For convenience, Table 1 summarizes the standardized latent variables

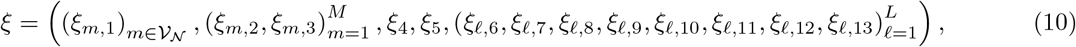

used in the MOFTy forward model with their corresponding interpretations in the model design. Here, *V*_*N*_ ⊆ *{*1, …, *M}* denotes the set of views with Gaussian likelihoods.

**Table 1.** Overview of the standardized latent variables used in the MOFTy forward model.

|  |  |
| --- | --- |
| $\xi_{m,1}$: | feature-wise noise precision variables for views with Gaussian likelihoods. |
| $\xi_{m,2}, \xi_{m,3}$: | Gaussian excitations scaled by inverse-gamma-transformed variables for view-specific weights. |
| $\xi_4$: | non-CFM components of the latent factors. |
| $\xi_5$: | factor-wise scaling variables to control the relative contribution of the CFM and non-CFM components. |
| $\xi_{\ell,6}$: | factor-specific CFM excitation field on the harmonic grid $\mathcal{G}^h$. |
| $\xi_{\ell,7}, \xi_{\ell,8}$: | zero mode and fluctuation variables of the factor-specific CFM amplitude spectrum. |
| $\xi_{\ell,9}, \xi_{\ell,10}$: | excitation variables for the integrated and direct Wiener components of the factor-specific power spectrum (one increment per grid step). |
| $\xi_{\ell,11}, \xi_{\ell,12}, \xi_{\ell,13}$: | variables controlling flexibility, asperity, and average slope of the factor-specific power spectrum. |

For views *m* ∈ *V*_*N*_, as in the classical MOFA model, we place a Gamma prior on the noise precision

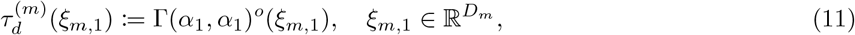

where Γ(*α*_1_, *α*_1_) denotes a Gamma distribution with shape and rate parameters both equal to *α*_1_ := 0.1, a small hyperparameter value often used to define a weakly informative Gamma prior. MOFA [11], MOFA+ [12], and MEFISTO [13] use two levels of sparsity regularization: automatic relevance determination (ARD [66]) to control factor relevance across views or groups, and spike-and-slab priors [67] to encourage sparsity of individual weights. By contrast, we use the following heavy-tailed shrinkage prior construction for the factor weights:

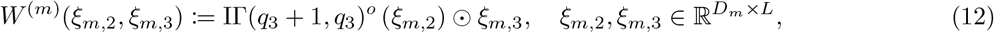

where ⊙ denotes element-wise multiplication and IΓ(*q*_3_ + 1, *q*_3_) denotes an inverse-gamma distribution with shape *q*_3_ + 1 and rate *q*_3_, with *q*_3_ := 0.1. Parameterizing shape and rate with this offset fixes the prior mean of the Gaussian scale at unity for any *q*_3_, leaving *q*_3_ to control tail heaviness alone: the induced marginal prior on each weight decays as 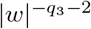, so small *q*_3_ gives a weakly informative, heavy-tailed prior. This construction can be interpreted as placing an inverse-gamma prior on the standard deviation of a zero-mean Gaussian scale mixture. Compared with placing an inverse-gamma prior on the variance, placing it directly on the standard deviation induces heavier tails and assigns substantially more prior mass to small Gaussian scales.

To see this, consider a single weight *w* = *σξ* with *ξ* ~ *N*(0, 1) and *σ* ~ IΓ(*a, b*) independent, corresponding to the parameterization above with *a* = *q*_3_ + 1 and *b* = *q*_3_. The marginal density is

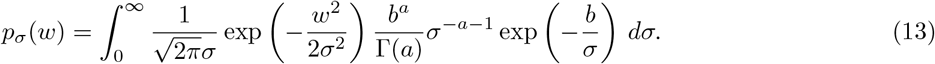

Substituting *σ* = |*w*|*u* factors out |*w*|^−*a*−1^ and leaves an integral that converges to a finite positive constant as |*w*| → ∞, so

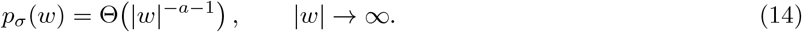

Under the usual variance mixture, *w* | *v* ~ *N*(0, *v*) with *v* ~ IΓ(*a, b*), the marginal density is

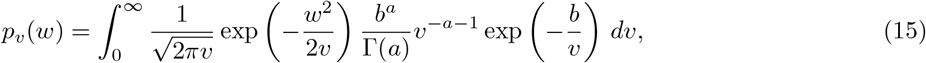

and the analogous substitution *v* = *w*^2^*u* gives

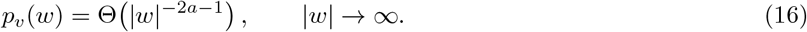

Since *a* + 1 *<* 2*a* + 1 for *a* > 0, the standard-deviation mixture has heavier polynomial tails at equal hyperparameters. The tail exponent alone, however, does not distinguish the two constructions, since it can be matched by retuning. If *v* ~ IΓ(*a*_*v*_, *b*_*v*_) and 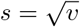, the induced density of the Gaussian scale *s* is

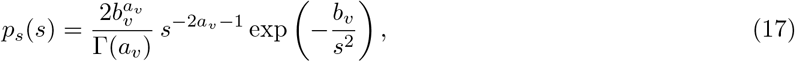

so the choice *a*_*v*_ = *a/*2 reproduces the polynomial factor *s*^−*a*−1^ of the direct standard-deviation prior exactly. The two priors then differ only in their exponential factors, exp(− *b*_*v*_*/s*^2^) against exp(− *b/s*), and this difference governs the behavior near zero. Let *Q*(*a, x*) = Γ(*a, x*)*/*Γ(*a*) denote the regularized upper incomplete gamma function, so that

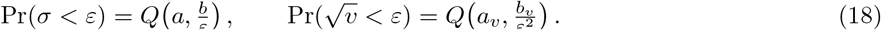

Since *Q*(*a, x*) ~ *x*^*a*−1^*e*^−*x*^*/*Γ(*a*) as *x* → ∞,

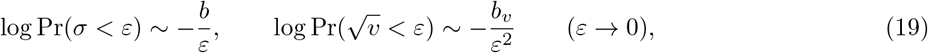

with the polynomial prefactors contributing only *O*(log(1*/ε*)). Hence 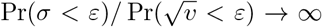 as *ε* → 0, for any values of the hyperparameters. An inverse-gamma prior on the variance can thus be retuned to match the polynomial tail behavior, but it cannot simultaneously reproduce the slower near-zero decay, and hence the greater prior mass assigned to very small Gaussian scales, of the direct standard-deviation prior. A further advantage of this formulation is that it avoids discrete latent variables, such as those introduced by spike-and-slab priors, which can be challenging for optimization. In MOFA-FLEX, this is addressed by providing horseshoe priors [68, 69] as an alternative to spike-and-slab priors.

In the absence of continuous covariates, we assume standard normal priors on the factors, i.e.,

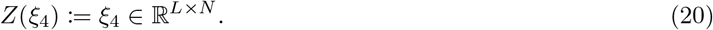

If the samples in our dataset are assumed to exhibit a correlation structure along continuous covariates *C* ⊆ ℝ^*C*^, we can incorporate this information in the prior structure with the nonparametric correlated field model in NIFTy [20, 21]. In this case, we define the forward model for the factors as the sum of a correlated field model (CFM) component and a non-CFM component

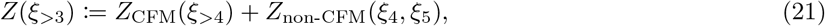

given by

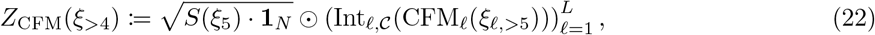

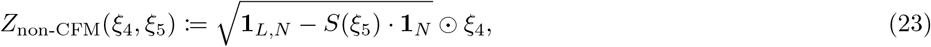

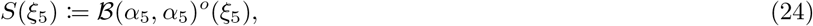

where *ξ*_4_ ∈ ℝ^*L*×*N*^, *ξ*_5_ ∈ ℝ^*L*^, **1**_*N*_ denotes an *N*-dimensional row vector of ones, **1**_*L,N*_ an *L × N* matrix of ones, and *B*(*α*_5_, *α*_5_) denotes the probability density of a Beta distribution with shape parameters both equal to *α*_5_ := 1. Note that the square-root operation is applied element-wise. For each factor *ℓ*, the correlated field model operator CFM_*ℓ*_ takes standardized latent variables *ξ*_*ℓ*,>5_ := (*ξ*_*ℓ*,6_, …, *ξ*_*ℓ*,13_) and transforms them into a Gaussian random field over a periodic grid space *G*. The construction of the correlated field model operator is provided in Section 4.3. For each factor, the corresponding correlated field is then evaluated at the covariates *C* using a factor-wise linear interpolation operator Int_*ℓ,C*_. The scaling operator *S* = *S*(*ξ*_5_) sets the mixing weights of the two components. Because the correlated field carries its own amplitude, the resulting prior variance ratio is *S ·* Var(CFM) : (1 − *S*) rather than *S* : (1 − *S*), so values of *S* close to one favor the CFM component more strongly than the nominal weight suggests. This formulation is conceptually related to the *ζ*_*ℓ*_ hyperparameter in MEFISTO and MOFA-FLEX, which controls the relative contribution of smooth and non-smooth components in the GP kernel.

### 4.3 Correlated field model

We briefly summarize the construction of the correlated field model as introduced in the original publication [20]. A more general formulation was later derived [21] and is also implemented in NIFTy. For additional background on information field theory, we refer the reader to its application to Bayesian field inference for local and causal dynamics [70].

To construct the grid space from a given set of continuous covariates *C* ⊆ ℝ^*C*^, we first determine the minimum spacing *h* in *C* across all coordinate directions.

We then scale and center *C* within a periodic Cartesian grid *G* with isotropic pixel size Δ*x*_*i*_ = *h* and *P*_*i*_ pixels per dimension. For the posterior approximations in this article, we applied padding that doubles the number of pixels in each coordinate direction, mitigating wrap-around effects arising from the periodic boundary conditions. In the implementation of MOFTy, Δ*x*_*i*_ is normalized to one by rescaling with *h*, which yields an equivalent representation. In practice, the grid was constructed by comparing different scalings and choosing one that adequately resolves the covariate domain (Section 4.7).

Since the a priori correlation structure is assumed to be homogeneous, the two-point correlation function depends only on the separation between two points and not on their absolute positions. By the Wiener–Khinchin theorem [26, 27], it can therefore be represented diagonally in harmonic space. Denote by *ℱ* : *G*^*h*^ → *G* the discrete Fourier transform and *G*^*h*^ the harmonic counterpart of *G* with pixel size 1*/*(*P*_*i*_Δ*x*_*i*_). For each factor *ℓ*, the correlated field model operator CFM_*ℓ*_ is defined as the product of an amplitude operator *A*(*ξ*_*ℓ*,>6_) and an excitation field *ξ*_*ℓ*,6_ on the harmonic grid, and the product is then transformed back to position space via the discrete Fourier transform, i.e.,

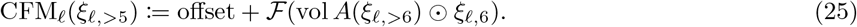

The volume factor vol = Π_*i*_ *P*_*i*_Δ*x*_*i*_ is chosen such that the zero mode equals the integral over position space, while the “offset” denotes the (known) mean of the field.

The log-normal operator LN(*m, s*) maps a standard normal random variable *ξ* ∈ ℝ to a log-normally distributed random variable with mean *m* > 0 and standard deviation *s* > 0:

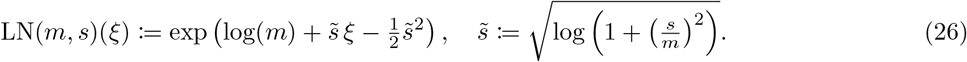

In NIFTy, the amplitude operator *A*(*ξ*_*ℓ*,>6_) is modeled as a spectral power field by combining a power-law component with an integrated Wiener process. The variable *ξ*_*ℓ*,7_ ∈ ℝ corresponds to the zero mode *A*_0_(*ξ*_*ℓ*,7_) = LN(*m*_7_, *s*_7_)(*ξ*_*ℓ*,7_). The nonzero modes are defined as

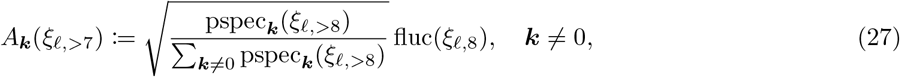

where fluc(*ξ*_*ℓ*,8_) := LN(*m*_8_, *s*_8_)(*ξ*_*ℓ*,8_) with *ξ*_*ℓ*,8_ ∈ ℝ models the amplitude of the field around its mean. The correlated field is further assumed to be isotropic, implying that the power spectrum depends only on the magnitude of the Fourier mode, |***k***|. For the nonzero modes ***k****≠* 0, the goal is to encode the following two prior assumptions in the model of pspec_***k***_:

1. field values are correlated along the given covariates *C*, favoring falling power spectra of the form pspec_|***k***|_ ~ _|_***k***| ^−*s*^ with *s* > 0;
2. sharp peaks in the power spectrum are unlikely and penalized.

These assumptions are incorporated by representing the power spectrum as

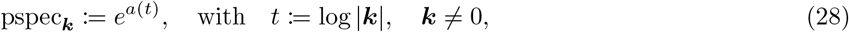

where *a*(*t*) is modeled as an integrated Wiener process, i.e.,

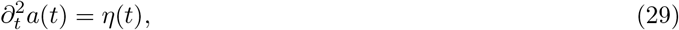

with Gaussian white noise *η*(*t*). To obtain a first-order representation suitable for forward-model evaluation, the derivative field *b*(*t*) of *a*(*t*) is introduced as an additional degree of freedom:

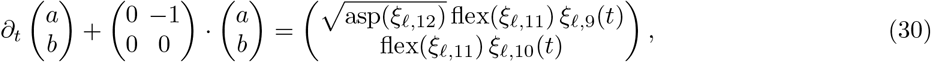

where flex(*ξ*_*ℓ*,11_) := LN(*m*_11_, *s*_11_)(*ξ*_*ℓ*,11_) (flexibility) and asp(*ξ*_*ℓ*,12_) := LN(*m*_12_, *s*_12_)(*ξ*_*ℓ*,12_) (asperity), with *ξ*_*ℓ*,11_, *ξ*_*ℓ*,12_ ∈ ℝ, and *ξ*_*ℓ*,9_(*t*), *ξ*_*ℓ*,10_(*t*) are independent standard Gaussian white-noise processes. Flexibility controls the amplitude of deviations from a power-law spectrum, whereas asperity controls the roughness of these deviations.

Solving Eq. 30 yields a Wiener process for the field *b*(*t*), driven by *ξ*_*ℓ*,10_. Consequently, *a*(*t*) is an integrated Wiener process for asp = 0, while for asp > 0, *ξ*_*ℓ*,9_ contributes an asperity-scaled, non-integrated Wiener-process component to *a*(*t*). The discrete representation of this solution, with initial condition 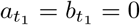, is given by

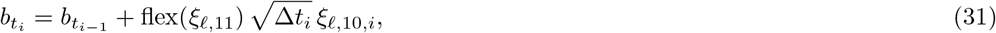

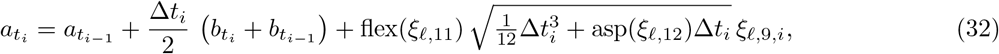

where *ξ*_*ℓ*,9_, *ξ*_*ℓ*,10_ collect the independent standard normal increments, one per step of the discretized grid. To control the average slope of the power spectrum, a further prior is introduced by replacing the endpoint difference of the integrated Wiener process with a linear trend term:

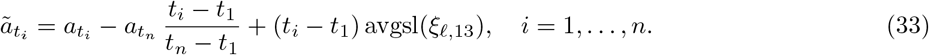

The average slope prior avgsl(*ξ*_*ℓ*,13_) := *s*_13_ *ξ*_*ℓ*,13_ + *m*_13_ is modeled as a normal random variable with mean *m*_13_ and standard deviation *s*_13_.

The CFM hyperparameters used in this article for posterior approximation and synthetic data generation with MOFTy are listed in Table 2 and can be flexibly adjusted. The current choice favors power spectra with negative slopes, so that white power tends to be attributed to the non-CFM component rather than absorbed into a flat CFM spectrum. This represents a prior preference rather than a constraint: where the data weakly determine the spectrum, near-flat spectra remain admissible. Conversely, correlated structure need not be absorbed by the CFM component. In the glioblastoma dataset (Section 2.2), for example, the posterior approximation of the non-CFM component exhibited positive autocorrelation despite this prior configuration.

**Table 2.** Hyperparameter settings of the NIFTy correlated field model for MOFTy posterior approximations and synthetic data generation. The numbers in brackets correspond to the indices of the latent variables *ξ*_*ℓ,i*_.

|  | Posterior approximation |  | Synthetic data generation |  |
| --- | --- | --- | --- | --- |
|  | mean | std | mean | std |
| Offset | 0 | — | 0 | — |
| [7] Zero mode | 2 | 1 | 1 | 0.5 |
| [8] Fluctuations | 2 | 1 | 3 | 0.5 |
| [11] Flexibility | 2 | 1 | 1 | 0.5 |
| [12] Asperity | 2 | 1 | 1 | 0.5 |
| [13] Average slope | -2 | 1 | -5 | 0.5 |

### 4.4 Inference and optimization

In this subsection, we summarize the main concepts behind metric Gaussian variational inference (MGVI [29]) and geometric variational inference (geoVI [28]). Terminology and notation follow the NIFTy documentation [71]. For a probability density *p*, we use the notation *H* := − log *p* for the information Hamiltonian, and denote expectation with respect to *p* by ⟨−⟩_*p*_.

Given the observed data *Y*, our goal is to approximate the posterior

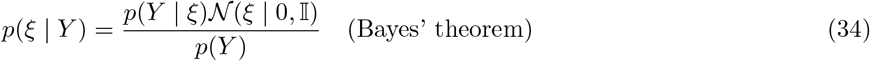

over the standardized variables *ξ*. Exact Bayesian inference in high-dimensional settings is generally intractable, because normalization, posterior summaries, and sampling all become computationally prohibitive. In such situations, variational inference (VI) provides a widely used approach for approximating *p* = *p*(*ξ* | *Y*) with a simpler parameterized family of distributions.

A common starting point is a Gaussian approximation

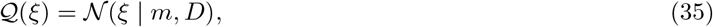

with mean *m* and covariance *D*. The parameters (*m, D*) of *Q* are determined by minimizing the variational Kullback–Leibler (KL) divergence between *Q* and the true posterior *p*:

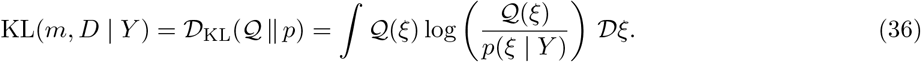

Since optimizing the full covariance is often infeasible, MGVI approximates the posterior precision *D*^−1^ by the Bayesian Fisher information metric [72, 73],

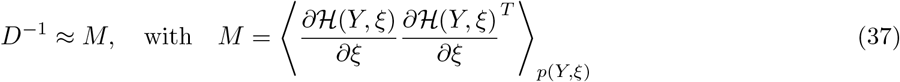

evaluated at the current mean *m*. In practice, this metric combines the curvature information contributed by the likelihood at *m* with the metric induced by the prior. As a consequence, the covariance is determined by this metric approximation, leaving only the mean *m* to be optimized variationally.

For fixed *D*, all terms in the KL divergence that do not depend on *m* can be dropped. The resulting objective is the posterior information Hamiltonian averaged under the current Gaussian approximation:

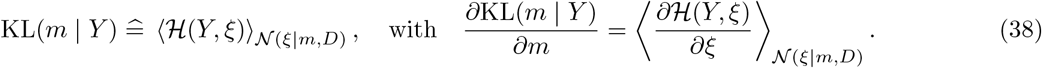

Both the KL divergence and its gradient are approximated using samples from the current variational distribution. Because the covariance is tied to a local metric approximation, MGVI tends to underestimate the posterior variance [29].

In settings where the posterior is nonlinear, a Gaussian approximation in the original standardized coordinates *ξ* may be inadequate. Geometric variational inference (geoVI) addresses this by constructing, around an expansion point 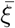, transformed coordinates 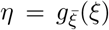 in which the posterior can be approximated by a standard Gaussian *Q*(*η*) = *N*(*η* | 0, I). This approximation is then pushed forward to the original coordinates through 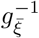:

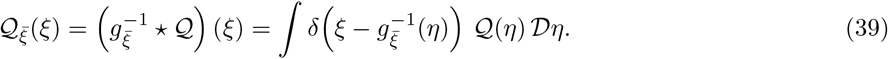

The expansion point 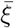 is chosen to minimize the KL divergence between 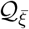 and the true posterior. In this way, geoVI captures curvature not only infinitesimally at the current mean, as in MGVI, but also in a local neighborhood around the optimal expansion point. Therefore, in highly nonlinear settings, geoVI can yield more accurate posterior approximations than MGVI, Gaussian full-covariance VI, and Gaussian mean-field VI, as demonstrated in the geoVI publication [28]. This makes geoVI an appropriate choice for MOFTy, where curved posterior geometries arise from several sources: the multiplicative coupling between factors and weights, the multiplicative coupling between the inferred power spectrum and the field excitations, and the heavy-tailed shrinkage priors on the weights.

During inference, MGVI or geoVI alternates between global iterations, where the variational distribution is updated, and internal iterations, where the posterior metric (MGVI and geoVI) and expansion points (geoVI) are optimized.

For the actual inference procedure, we combined geoVI with maximum a posteriori (MAP) estimates. Empirically, this combination improved numerical stability during training. MAP estimates were used for the noise model variables *ξ*_*m*,1_. For the factor weights, MAP estimates were used for the inverse-gamma scale component of the prior (*ξ*_*m*,2_), while posterior samples were retained for the weights *W* ^(*m*)^ themselves. In the correlated field component of each factor, MAP estimates were used for the power spectrum variables *ξ*_*ℓ*,7_, *ξ*_*ℓ*,8_, *ξ*_*ℓ*,11_, *ξ*_*ℓ*,12_, and *ξ*_*ℓ*,13_, corresponding to the zero mode, fluctuations, flexibility, asperity, and average slope, respectively, as well as for the scaling variables *ξ*_5_. Posterior samples were retained for the integrated Wiener process increments *ξ*_*ℓ*,9_ and *ξ*_*ℓ*,10_, which carry the deviations of the log-log power spectrum from its power-law trend and therefore its nonparametric shape.

The following optimization settings, used for all posterior approximations in this article, follow established NIFTy best practices. For all models in this article, we ran 30 global iterations in total. Each global iteration consists of conjugate gradient linear sampling steps for inverting the posterior metric (MGVI) and additional nonlinear sampling steps for geoVI, followed by KL optimization. The optimization parameters are summarized in Table 3. At each iteration, we configured the algorithm to draw four antithetic mirrored sample pairs, yielding eight samples in total. Each model in this article was trained with four random seeds, and after the last global iteration, posterior samples were summarized across seeds as described in Section 4.5. Thus, we obtained 32 posterior samples in total, which were used to compute posterior means and uncertainty estimates.

**Table 3.** Optimization parameters for each global iteration (GI) with number of iterations I(−), minimizer method (conjugate gradient (CG), vector-free limited-memory BFGS [74] (VL-BFGS)), convergence criterion (Δ*E*), and convergence level (Level).

| Total GIs : 30 | $I(\text{GI} \leq 5)$ | $I(\text{GI} > 5)$ | Method | $\Delta E$ | Level |
| --- | --- | --- | --- | --- | --- |
| Linear sampling | 200 | 1000 | CG | 0.5 | 1 |
| Nonlinear sampling | 50 | 300 | VL-BFGS | 0.5 | 2 |
| KL optimization | 200 | 1000 | VL-BFGS | 0.5 | 2 |

**Table 4.** Training performance of MOFTy and MEFISTO. Peak memory is reported as maximum resident set size (CPU) and maximum allocated GPU memory, reported as approximate total and per-model values. MOFTy models were trained concurrently with four random seeds; MEFISTO was trained with a single seed, so per-model values provide a direct comparison. Runtime is wall-clock time for the full training run, measured in hours (h). *N* = 5,573 denotes the number of spatial observations and *U* the number of inducing points.

| Method | Dataset and configuration | System | Peak memory, total (per model) |  | Runtime |
| --- | --- | --- | --- | --- | --- |
|  |  |  | CPU | GPU |  |
| MEFISTO | Glioblastoma, full GP ($U = N$) | 2 (GPU) | 13.8 GB (13.8 GB) | 781 MB (781 MB) | 9.3 h |
| MEFISTO | Glioblastoma, $U = 3,000$ | 2 (GPU) | 6.7 GB (6.7 GB) | 781 MB (781 MB) | 3.5 h |
| MEFISTO | Glioblastoma, $U = 1,000$ | 2 (GPU) | 4.6 GB (4.6 GB) | 781 MB (781 MB) | 0.4 h |
| MOFTy | Glioblastoma, ten factors | 2 (GPU) | 3.2 GB (800 MB) | 13.7 GB (3.4 GB) | 24.3 h |
| MOFTy | Glioblastoma, eight factors | 2 (GPU) | 3.2 GB (800 MB) | 13.3 GB (3.3 GB) | 19.9 h |
| MOFTy | Glioblastoma, six factors | 2 (GPU) | 3.2 GB (800 MB) | 12.5 GB (3.1 GB) | 15.3 h |
| MOFTy | Glioblastoma, four factors | 2 (GPU) | 2.8 GB (700 MB) | 12.2 GB (3.1 GB) | 11.7 h |
| MOFTy | Glioblastoma, four factors | 1 (CPU) | 5.3 GB (1.3 GB) | — | 21.8 h |
| MOFTy | Mouse gastrulation | 1 (CPU) | 3.7 GB (925 MB) | — | 8.1 h |
| MOFTy | 1D synthetic (900 points) | 1 (CPU) | 2.6 GB (650 MB) | — | 1.9 h |
| MOFTy | 2D synthetic ($30 \times 30$ pixels) | 1 (CPU) | 2.3 GB (575 MB) | — | 1.8 h |
| MOFTy | 2D synthetic ($100 \times 100$ pixels) | 2 (GPU) | 3.2 GB (800 MB) | 29.7 GB (7.4 GB) | 27.8 h |

For all posterior approximations in this article, we initialized *Z*_non-CFM_ using PCA scores computed from the concatenated preprocessed data matrices across all views. For this PCA initialization, missing values, when present, were imputed using the feature-wise mean. We further initialized *ξ*_5_ such that *S*(*ξ*_5_) ≈ 0, allowing covariate-dependent structure to emerge gradually during optimization. Together with PCA-based initialization, this provides a data-driven initial orientation of the latent space and reduces sensitivity to arbitrary random initializations during optimization. The remaining trainable parameters were initialized by drawing from a standard normal distribution and multiplying by 0.1 to reduce numerical instabilities at the beginning of training. To prevent overfitting during the initial training phase, we fixed the noise precision for Gaussian likelihoods at 1.0 in iterations 1 to 5, and inferred it jointly with all remaining parameters in iterations 6 to 30.

### 4.5 Posterior alignment and uncertainty estimation

The operator-based implementation in NIFTy allows us to generate samples at each layer of the forward model by applying the corresponding operators to posterior samples of the standardized variables *ξ*.

After training, we pooled posterior samples obtained from different random seeds. In the absence of the CFM component, the likelihood together with the isotropic latent Gaussian priors would retain the usual non-identifiability of factor models under orthogonal transformations. The full model is generally not invariant under arbitrary orthogonal transformations, because the CFM component assigns factor-specific power spectra to model the correlation structures. Mixing factors can therefore introduce cross-spectra between the CFM components of different factors. We nonetheless used orthogonal alignment because it preserves distances, angles, and overall scale in the latent factor space, avoiding arbitrary rescaling or shearing of the factors and weights.

We selected one seed as a reference and aligned the posterior mean of the factors from all other seeds to the reference using orthogonal Procrustes analysis [75, 76], yielding seed-specific orthogonal transformation matrices. For non-negative factorizations, the non-negativity constraint already removes the rotational freedom, leaving only a permutation ambiguity; MOFTy therefore aligns factors across seeds using linear sum assignment with Pearson correlation as the cost function. The seed-specific orthogonal transformations were then applied to the posterior samples of the weights and factors, as well as to the posterior samples of the CFM component and the non-CFM component. For the factor-wise power spectra, we applied the element-wise square of the transformation matrix to the posterior samples of the factor-wise spectra, which yields an approximation to the exact transformation of the spectra under orthogonal mixing.

To evaluate recovery on the synthetic data, where the ground-truth factors are known, we aligned the posterior mean of the factors from each seed directly to the ground truth, so that recovery is assessed up to an orthogonal transformation.

Finally, we concatenated the aligned posterior samples across seeds. From these pooled samples we computed posterior means and uncertainty estimates for the weights, the combined factors and their CFM and non-CFM components, and the factor-wise power spectra.

### 4.6 Performance and computational complexity

For the correlated field model, the most computationally demanding step is the fast Fourier transform (FFT). Because one FFT is required per factor, the FFT contribution to a single likelihood evaluation scales as *O*(*LP* log *P*), where *L* denotes the number of latent factors and *P* the number of grid points in harmonic space. This contrasts with sparse inducing-point GP approximations, for which the dominant cost scales as *O*(*LNU* ^2^) with *N* observations and *U* ≤ *N* inducing points. The total cost of a likelihood evaluation in the full factor model also includes interpolation and matrix multiplications between factors and weights. Computation of the sampled KL divergence requires evaluating the likelihood once per sample. Memory consumption scales linearly with the number of samples used for the KL approximation and the size of the inference problem. This linear scaling is enabled by the implicit operator-based implementation in NIFTy, which represents the posterior metric implicitly and avoids explicitly storing large matrices during inference. Although the computational cost of an individual KL evaluation scales near-linearly with problem size, the overall algorithm may scale superlinearly, since larger problems typically require more optimization steps to converge.

#### 4.6.1 Implementation

For larger datasets, NIFTy additionally provides GPU acceleration via CuPy [77] and parallelized sampling using MPI [78]. Future versions of MOFTy could also be implemented using the JAX [79] version of NIFTy [25], enabling additional acceleration. In contrast, the current implementation uses the NumPy [80]-based implementation of NIFTy, which emphasizes modular design with minimal external dependencies.

#### 4.6.2 Training performance

We use the following abbreviations: GB (gigabyte), MB (megabyte), and TB (terabyte). Total training time was measured as wall-clock time and is reported in hours per dataset. Two systems were used to train MOFTy and MEFISTO [13] models:

- **System 1:** An Apple MacBook Pro (14-inch, November 2024) running macOS 15.1 (24B2083), equipped with an Apple M4 Max chip and 36 GB of unified memory.
- **System 2:** A Linux workstation running Ubuntu 24.04.4 LTS, equipped with two 64-core Intel Xeon Gold 6430 processors (two threads per core), 1 TB of main memory, and two NVIDIA A100 GPUs with 80 GB of memory each.

Peak memory usage was measured as maximum resident set size and, on System 2, additionally as maximum allocated GPU memory. It is reported as a dataset total and as an approximate per-model (seed) value.

On System 1, eight MOFTy models were trained in parallel, each on a single CPU core. Specifically, the MOFTy models for the glioblastoma (four factors) and mouse gastrulation datasets were trained concurrently using four random seeds each, yielding eight parallel models in total. Likewise, the 1D and 2D synthetic data reconstructions (Fig. 1b–d) were trained concurrently with four random seeds each.

On System 2, we trained four models (four random seeds) on a single GPU for the glioblastoma factor robustness analysis and the large 2D synthetic data reconstruction (Ext. Data Fig. 4). The four-factor glioblastoma models were additionally trained on a single GPU for performance comparison, while the CPU-trained models were used for all downstream analyses. For the present datasets, we observed that training more than four models concurrently on System 2 substantially increased total training time, likely due to GPU/CPU memory contention and increased data-transfer overhead.

For the comparison analysis on the glioblastoma dataset, MEFISTO models were trained in GPU mode with four latent factors, one random seed (42), convergence mode “slow”, 16 MKL threads, and three model configurations: 1,000 and 3,000 inducing points, as well as a full GP approximation (5,573 spatial spots). All models were trained separately on one GPU.

### 4.7 Preprocessing

Preprocessing was performed using the MOFA2 R package and its Python interface mofax [11–13], together with muon [31], scanpy [36], and squidpy [81]. The preprocessed datasets were then stored in a MOFA2- and muon-compatible HDF5 format, which was subsequently used for training with MOFTy.

#### 4.7.1 Glioblastoma dataset

The publicly available glioblastoma dataset was obtained from 10x Genomics (Code and data availability, Section 5). Preprocessing was performed with muon, scanpy, and squidpy. For the gene expression modality, we removed spots below the first percentile and above the 99th percentile of total counts, and spots below the first percentile of detected genes. Mitochondrial transcript abundance was not used for spot-level filtering, as elevated mitochondrial signal in spatial spots may reflect biologically meaningful variation, including metabolic heterogeneity and hypoxic regions, and is therefore not necessarily indicative of low-quality measurements [82]. Among 35 antibody features, four isotype controls (mouse IgG2a, mouse IgG1k, mouse IgG2bk, rat IgG2a) were excluded, and only the 31 biological proteins were retained. Protein-based filtering (using biological proteins) removed spots below the first percentile of total protein counts and below the first percentile of detected proteins. We did not exclude spots above the 99th percentile of total protein counts because, for a small targeted protein panel, extreme total protein counts are not necessarily technical outliers and may reflect genuine biological enrichment. After spot filtering, the dataset contained 5,573 spatial spots, compared with 5,756 spots under tissue in the raw dataset. Gene features were retained if they were detected in at least 10 of the 5,573 filtered spots. Biological protein features were filtered using the same threshold, which did not remove any protein features in this dataset. For the CFM configuration, we mapped the spatial spot coordinates onto a regular grid with 94 × 110 pixels, on which the area covered by the spots amounted to 5,620 pixels. We added grid padding in each direction, resulting in a total grid size of 188 × 220.

Normalization was performed separately by modality. Gene expression counts were library-size normalized and transformed as log(1 + *x*), and the top 2,000 highly variable genes were retained. Values of the biological protein features were transformed using a spot-wise centered log-ratio (CLR) transformation. Next, we standardized each modality: each feature was centered, and each modality was scaled by its overall standard deviation. MOFTy was then initialized with four, six, eight, or ten latent factors, using Gaussian likelihoods for both modalities.

#### 4.7.2 Mouse gastrulation dataset

The preprocessed dataset, including the 2D UMAP coordinates [47, 48], was obtained from the corresponding MEFISTO tutorial [30]. As is typical for neighbor-based embeddings such as UMAP, the 2D UMAP coordinates show substantial variation in local point density. For the CFM configuration, we therefore mapped the UMAP coordinates onto a 53 × 65 pixel grid; the area enclosed by the data points covered 912 of these pixels. The CFM component is therefore band-limited below the data resolution by construction. We then added padding in each direction, resulting in a final grid size of 106 × 130 pixels.

Before training, we standardized each modality: each feature was centered, and each modality was scaled by its overall standard deviation. We then initialized MOFTy with ten factors, matching the MEFISTO publication [13], and used Gaussian likelihoods for all three modalities.

### 4.8 Synthetic data generation

For the simulations, we generated synthetic data for three views with the MOFTy forward model by drawing one *ξ* sample for each operator (random seed 42), assuming a CFM prior and a Gaussian likelihood without non-negativity constraints on the factors or weights. The CFM hyperparameters are listed in Table 2 (synthetic data generation). For the one-dimensional case, we generated five latent factors, 900 points on a one-dimensional periodic grid (no padding), and 300 features per view (three views in total). For the two-dimensional case, we generated five latent factors, 900 points on a 30 × 30 periodic grid (no padding), and 300 features per view (three views in total). To assess scalability, we also generated synthetic data with five latent factors, 10,000 samples on a 100 × 100 periodic grid (no padding), and 1,000 features per view (three views in total). We multiplied each sampled excitation *ξ*_*i*_ by 0.6. The factor-wise variance scaling operators were set to 0.99, 0.75, 0.5, 0.25, and 0.01 for factors 1–5, respectively. Because the correlated field carries its own amplitude (prior mean 3.0 for the fluctuations; Table 2, synthetic data generation), these mixing weights correspond to CFM variance shares of approximately 0.999, 0.96, 0.90, 0.75, and 0.08. In this way, we obtained latent factor components *Z*_CFM_ and *Z*_non-CFM_, their sum *Z* = *Z*_CFM_ +*Z*_non-CFM_, and view-specific weights *W* ^(*m*)^. Together, these define the noiseless synthetic signal *s*^(*m*)^ = *W* ^(*m*)^*Z*. We then generated feature-wise additive Gaussian noise *ϵ*^(*m*)^, with precisions sampled from the Gamma prior, and scaled it such that the signal-to-noise ratio 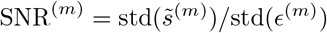 equaled one in each view *m*, where 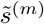 denotes the synthetic signal centered feature-wise across samples. The synthetic data were then obtained via *Y* ^(*m*)^ = *s*^(*m*)^ + *ϵ*^(*m*)^ and 20% of the data columns were randomly masked in each view to simulate missing data.

For the posterior approximations, we initialized MOFTy with five latent factors and standardized each view before training: each feature was centered, and each view was scaled by its overall standard deviation. We used Gaussian likelihoods for all views and the same model configurations as for the glioblastoma and mouse gastrulation datasets, including the CFM hyperparameters used for posterior approximation (Table 2), and PCA initialization of the non-CFM component. We also used the same grid padding in each coordinate direction, resulting in a grid of 1,800 points for the one-dimensional case and grids of 60 × 60 and 200 × 200 pixels for the two-dimensional cases, respectively.

## 5 Code and data availability

- MOFTy is a Python package distributed under the GNU General Public License v3.0 and available at: https://github.com/neumann-mn/mofty. It includes the implementation of MOFTy and all scripts to reproduce the results in this article.
- NIFTy is distributed under the GNU General Public License v3.0 and available at: https://gitlab.mpcdf.mpg.de/ift/nifty.
- The human glioblastoma spatial dual-omics dataset was obtained from 10x Genomics and is available at: https://www.10xgenomics.com/datasets/gene-and-protein-expression-library-of-human-glioblastoma-cytassist-ffpe-2-standard.
- The mouse gastrulation dataset is available as part of the original publication [33] and the MEFISTO publication [13]. The version used in this article can be accessed from the corresponding MEFISTO tutorial [30].

## 6 Acknowledgments

- M.N. was supported by the Helmholtz Association under the joint research school “HIDSS4Health – Helmholtz Information and Data Science School for Health”.
- M.N. and A.O. were supported by the Karlsruhe Institute of Technology and the Helmholtz Association [POF4; 5207.0004.0012].
- This project was supported by the Vector Stiftung.

## 7 Author contributions

- M.N. conceived the project and designed the method.
- M.N. performed all analyses.
- M.N. implemented the software package and is its lead developer and maintainer.
- M.N. wrote the manuscript and generated all figures.
- P.A. provided advice on (numerical) information field theory (IFT/ NIFTy) and gave methodological guidance throughout the project.
- P.A., A.K.K, and A.O. critically revised the manuscript.
- A.K.K. and A.O. supervised the project and provided project funding and computational resources.

## 9 Extended Data Figures

**Extended Data Fig. 4.**
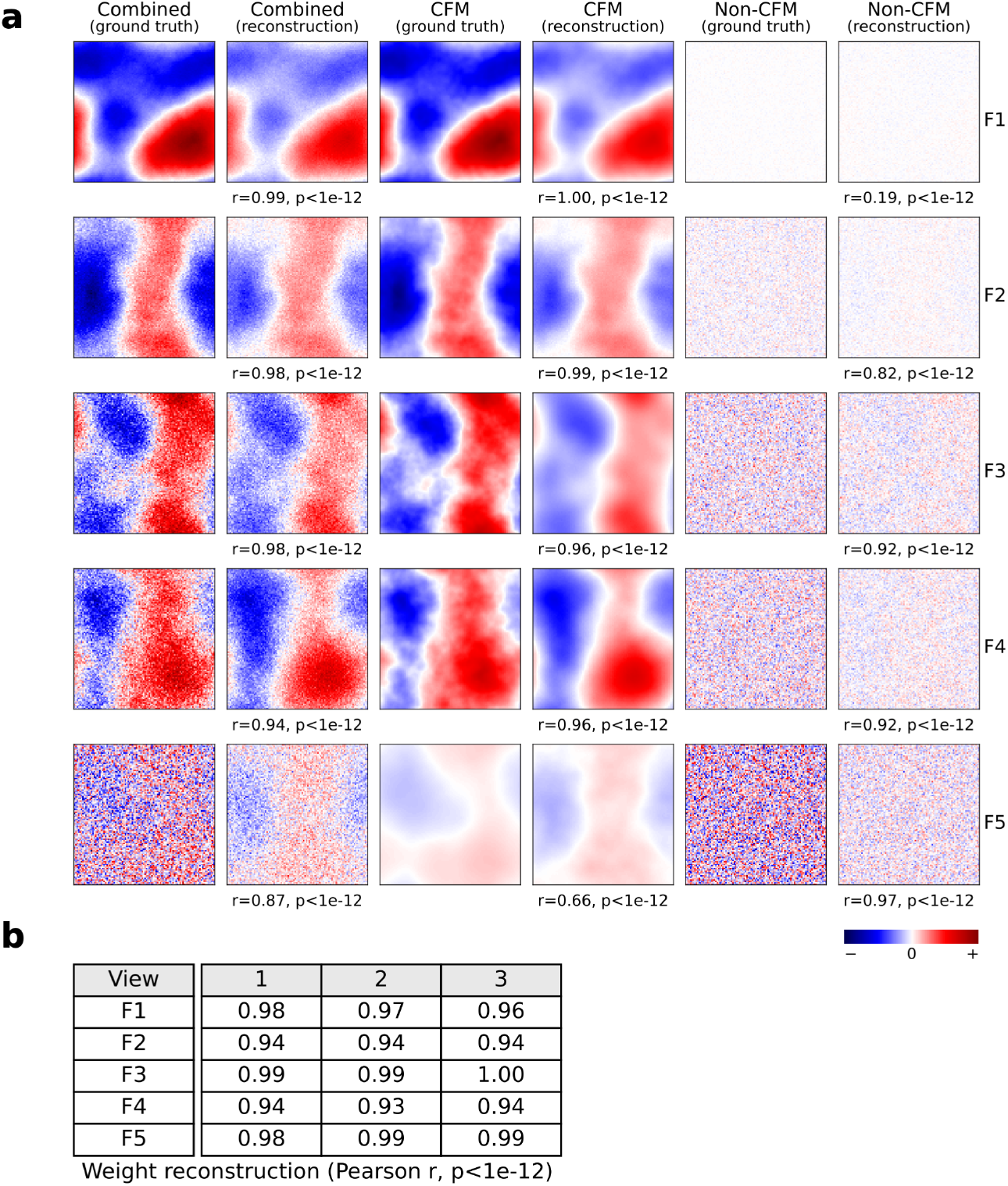
**a**, Synthetic trimodal dataset with 2D covariates (100*×*100 pixels) and known ground truth (five factors, F1–F5). Pearson correlation coefficients (*r*) with two-sided *p*-values between the ground truth and inferred weights/factors are reported. Heatmaps display the factor values. Values of the combined, CFM, and non-CFM components are displayed at spatial coordinates using a shared normalized color scale across components. **b**, Inferred weights are compared to the ground truth weights.

**Extended Data Fig. 5.**
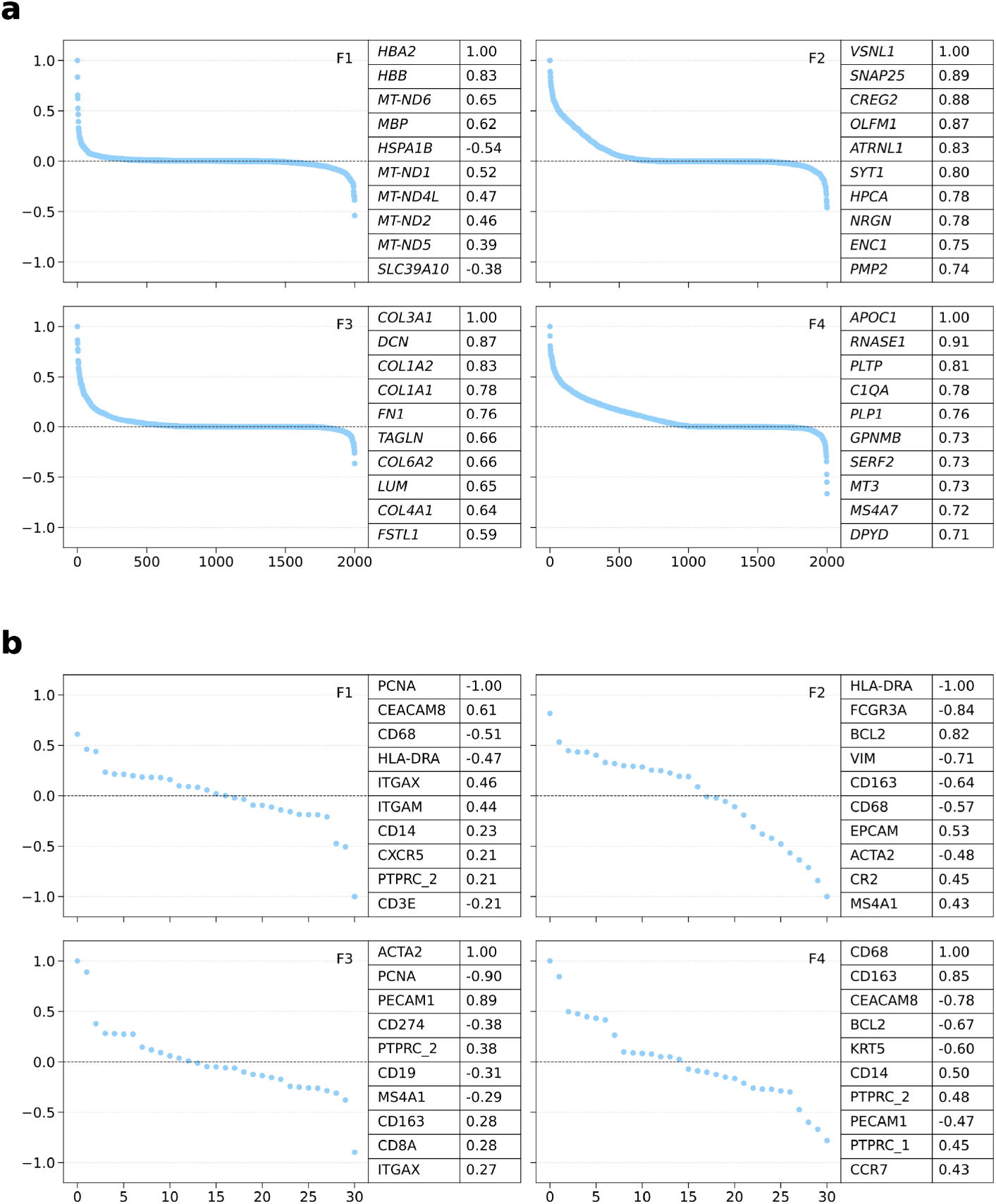
Visualization of gene expression (**a**) and protein expression (**b**) weights in the glioblastoma dataset for factors 1–4. The tables show the top 10 features ranked by absolute weight for each factor, together with the sign of the corresponding weight. The weights are normalized by the maximum absolute weight value within each factor.

**Extended Data Fig. 6.**
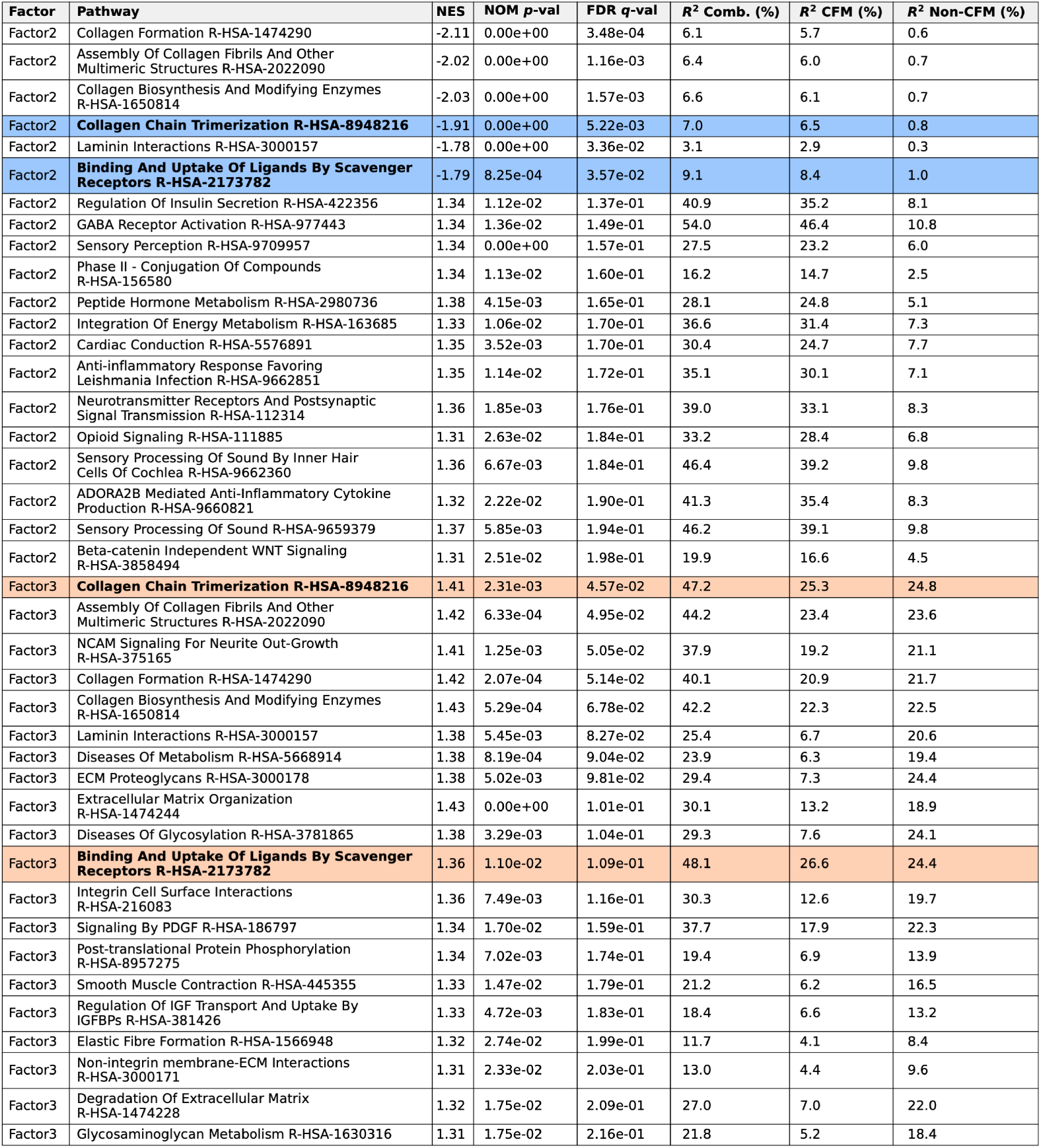
Gene set enrichment analysis (GSEA) of the gene expression modality weights for factors 2 and 3 in the glioblastoma dataset. For each factor, the tables show the top 20 enriched pathways ranked by FDR-adjusted *q*-value (ascending), NOM *p*-value (ascending), and NES (descending by absolute value) using GSEApy [37] and Reactome [38]. For each factor, we only included pathways with an FDR-adjusted *q*-value below 0.25 and a nominal *p*-value below 0.05. Variance explained (*R*^2^) by the specified factor components in the corresponding gene sets is reported as percentages. The sign of the normalized enrichment score (NES) indicates the direction of association with the factor, i.e., enrichment among features with negative or positive factor weights. The pathways shown in Fig. 2c are highlighted. Here, “Collagen chain trimerization” and “Scavenger receptor uptake” denote shortened labels for “Collagen Chain Trimerization R-HSA-8948216” and “Binding And Uptake Of Ligands By Scavenger Receptors R-HSA-2173782”, respectively.

**Extended Data Fig. 7.**
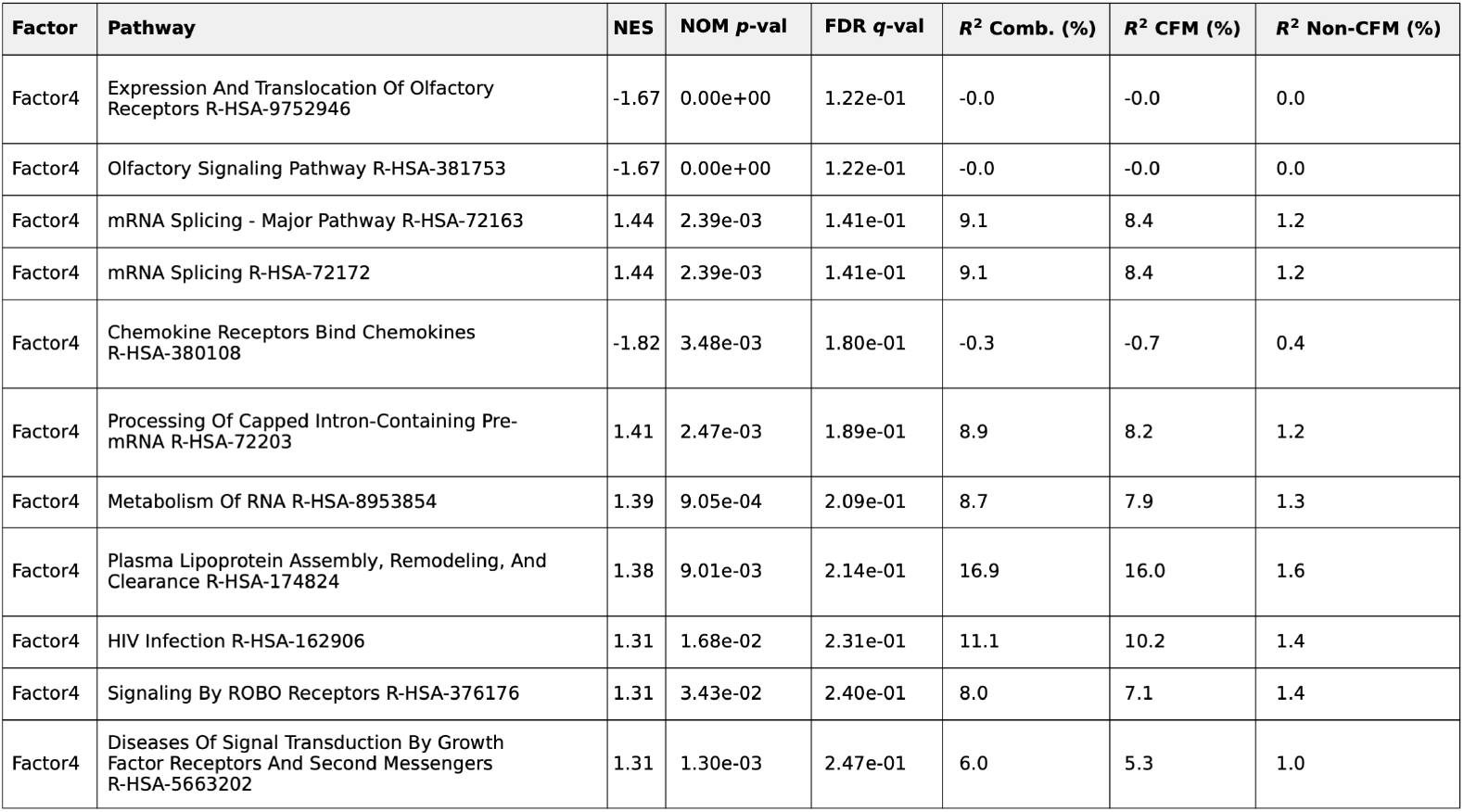
Gene set enrichment analysis (GSEA) of the gene expression modality weights for factor 4 in the glioblastoma dataset. The analysis is analogous to that in Ext. Data Fig. 6.

**Extended Data Fig. 8.**
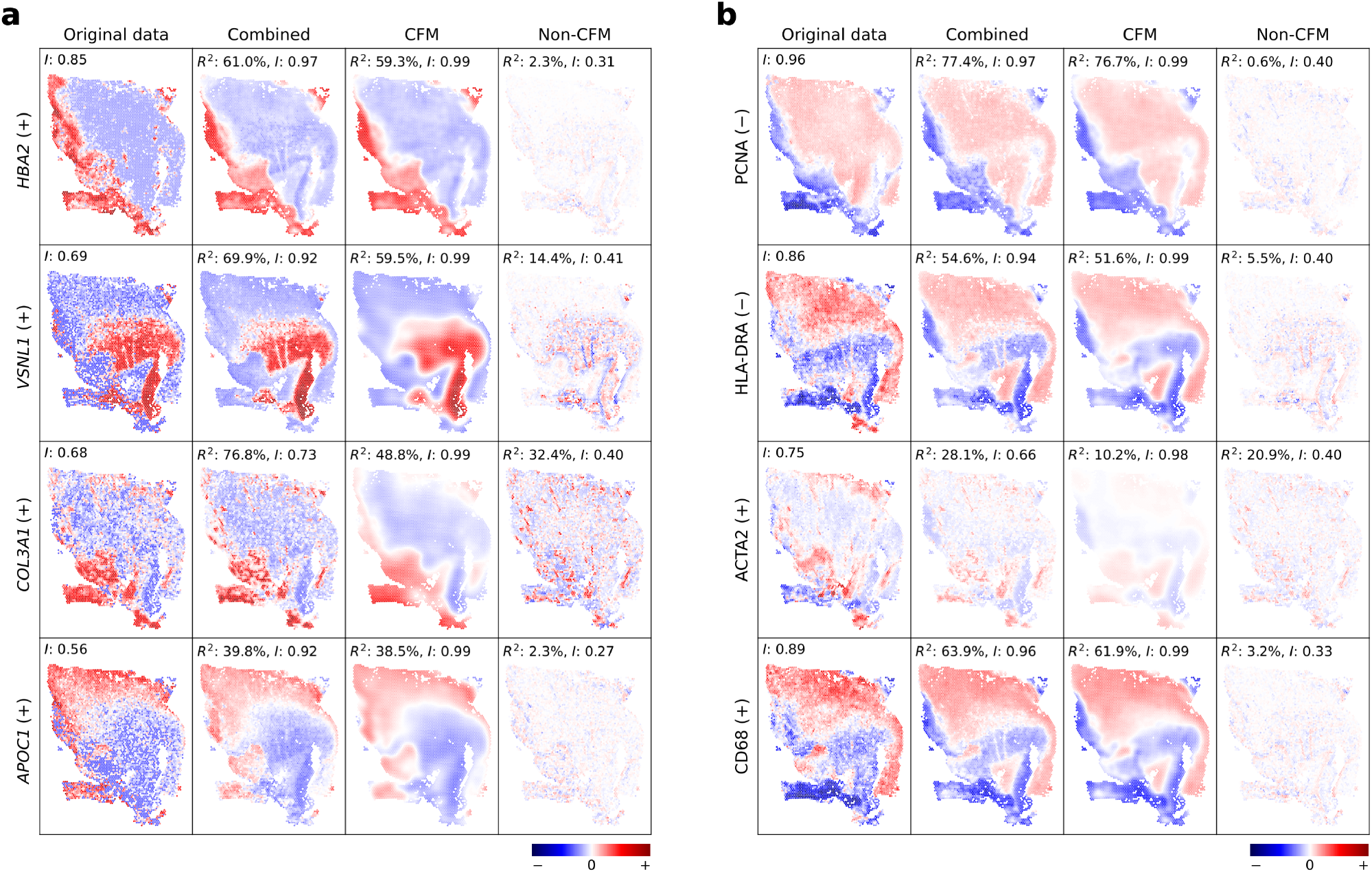
Feature reconstruction in the glioblastoma dataset for the features with the highest absolute weights in gene expression (**a**) and protein expression (**b**) modalities for factors 1–4 (top to bottom). The sign of the corresponding feature weight is indicated in brackets. The reconstruction for each feature was obtained by multiplying the corresponding feature weight vector by the combined, CFM, or non-CFM factor matrix. Each row was plotted on the same normalized color scale.

**Extended Data Fig. 9.**
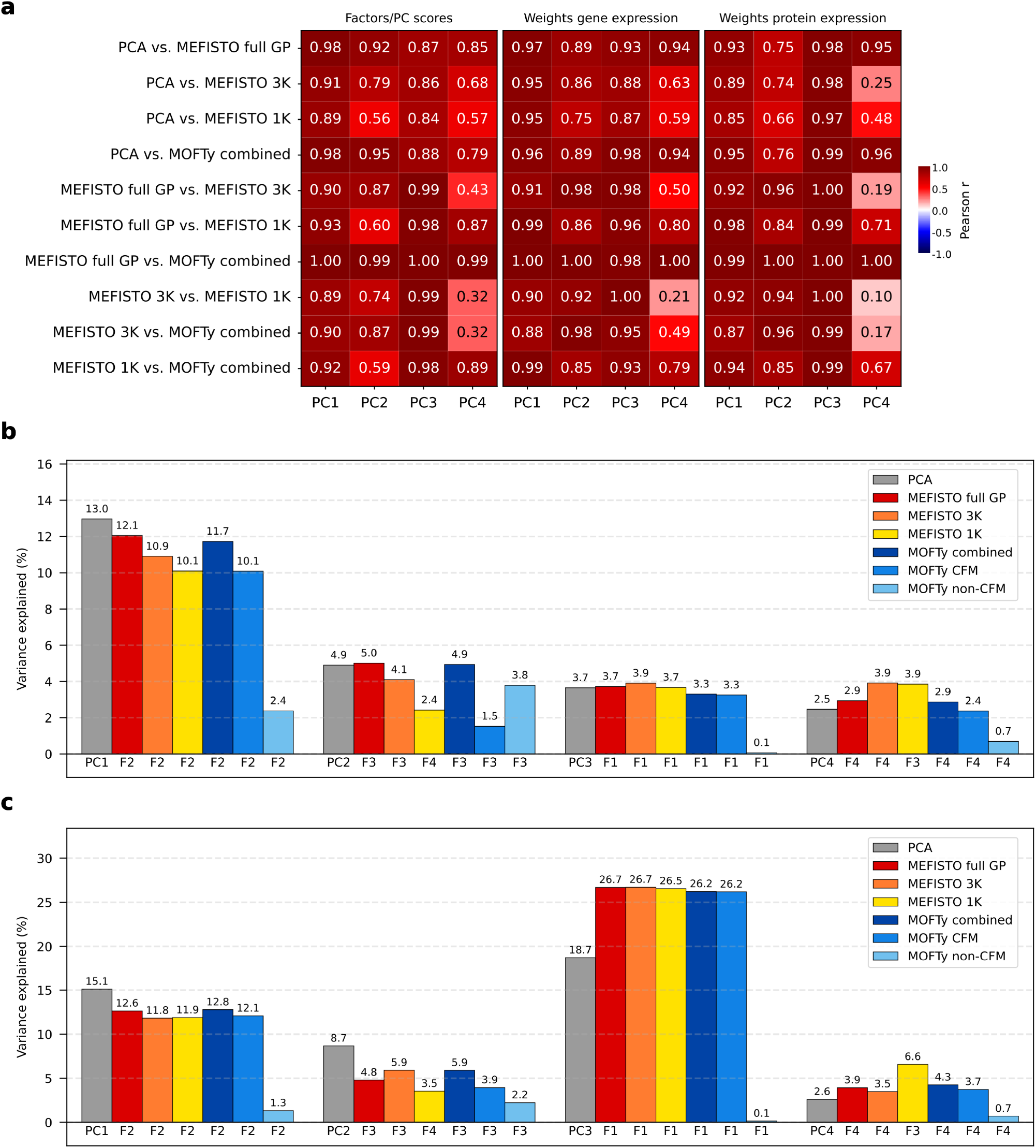
Comparison of MEFISTO, PCA, and MOFTy on the glioblastoma dataset. **a**, We used PCA scores as anchors and matched the factors of MEFISTO with 1,000 inducing points, MEFISTO with 3,000 inducing points, and MEFISTO with a full GP approximation, as well as the MOFTy combined factors to the PCA scores using linear sum assignment with absolute Pearson correlation as the cost function. Shown are the Pearson correlation coefficients between all models and the matched factors according to the PCA anchors. These assignments are consistent with visual inspection of the factor values over the spatial spots in Ext. Data Fig. 10. **b, c**, Shown are the variance explained (*R*^2^) values of the matched PCA scores and the factors of the MEFISTO models and MOFTy for the gene expression (**b**) and protein expression (**c**) modalities.

**Extended Data Fig. 10.**
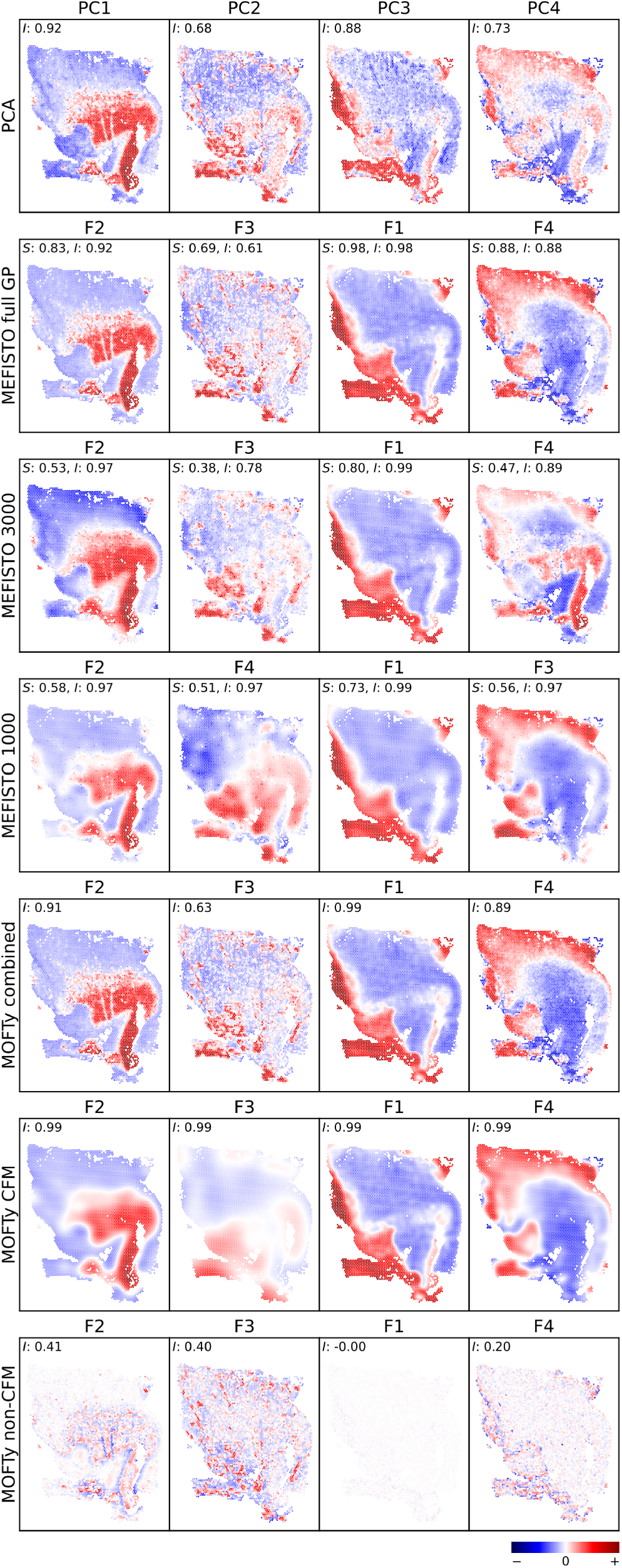
Spatial comparison of MEFISTO, PCA, and MOFTy factors in the glioblastoma dataset. Displayed are scores over the spatial spots for the first four principal components from PCA [34, 35] in the first row. MEFISTO with 1,000 inducing points is shown in the second row, and MEFISTO with 3,000 inducing points is shown in the third row. In the fourth row, MEFISTO with a full GP approximation is shown. The combined MOFTy factors and their CFM and non-CFM components from the four-factor configuration are shown in the fifth, sixth, and seventh rows, respectively, plotted factor-wise on the same normalized color scale. MEFISTO factor values and PCA scores are also displayed using a shared normalized color scale. For each panel, Moran’s *I* is reported, reflecting the degree of spatial autocorrelation in the corresponding factor values. For the MEFISTO models, the inferred GP variance-scale parameter *S* is shown for each factor, giving the fraction of factor variance attributed to the smooth GP component. All columns were ordered according to the assignment based on Ext. Data Fig. 9a.

**Extended Data Fig. 11.**
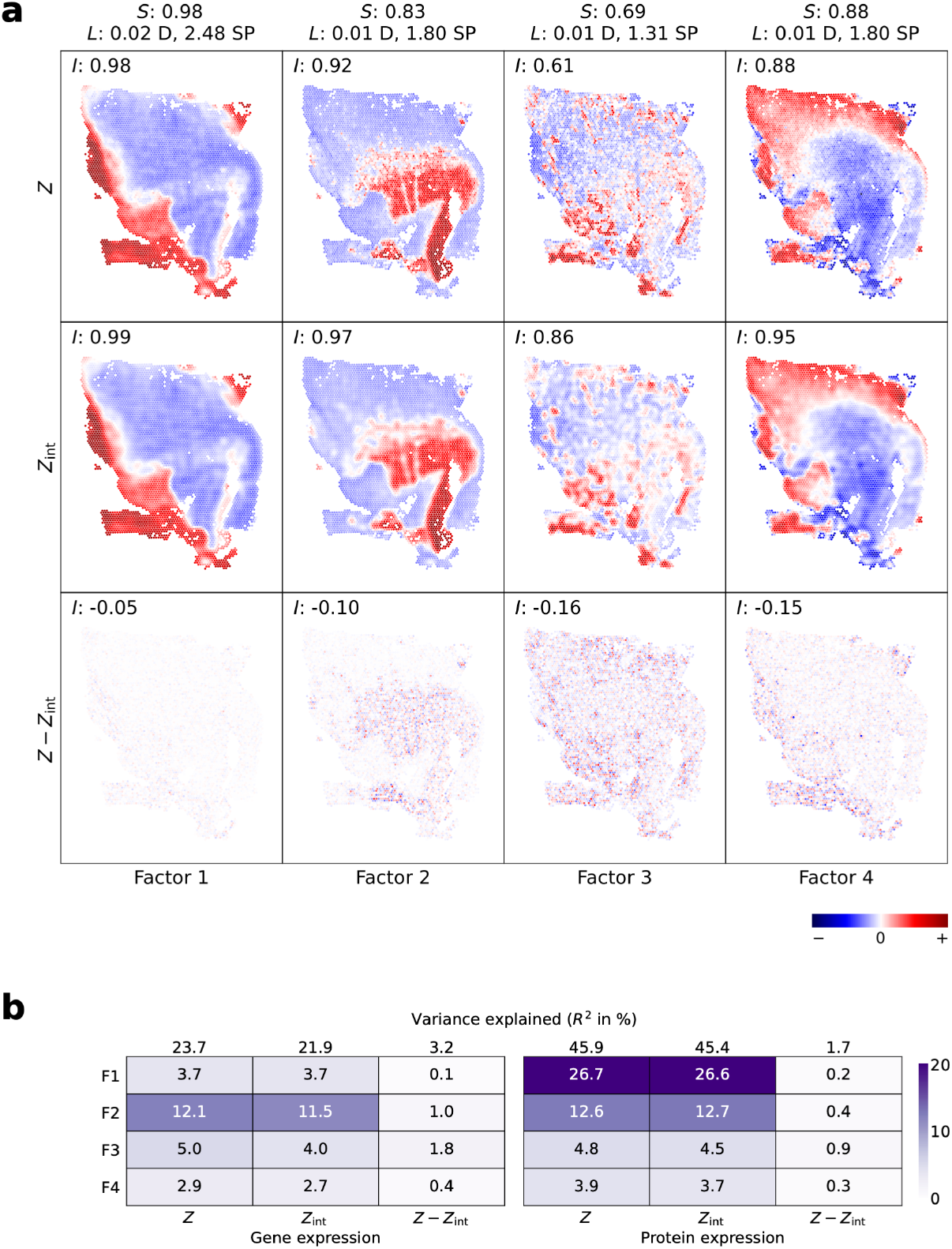
MEFISTO factor decomposition on the glioblastoma dataset. **a**, Spatial maps of the four MEFISTO factors (columns), showing the factor values (*Z*), their interpolation at the observed coordinates (*Z*_int_), and the difference (*Z* − *Z*_int_). Factor values are color-coded after scaling each factor by the maximum absolute value across the three components. Column headers show the learned variance-scale parameter *S* and length scale *L*; the length scale is reported both in units of the spatial domain size (D, the extent of the covariate range) and in units of the median nearest-neighbor spot spacing (SP). Each panel is annotated with Moran’s *I*, quantifying spatial autocorrelation. **b**, Variance explained per factor for each component, shown separately for the gene expression and protein expression modalities. Total variance explained per modality is shown above each heatmap. Because the components are propagated through the shared loading matrix and are not statistically independent, their contributions are not additive and do not sum to the combined total.

**Extended Data Fig. 12.**
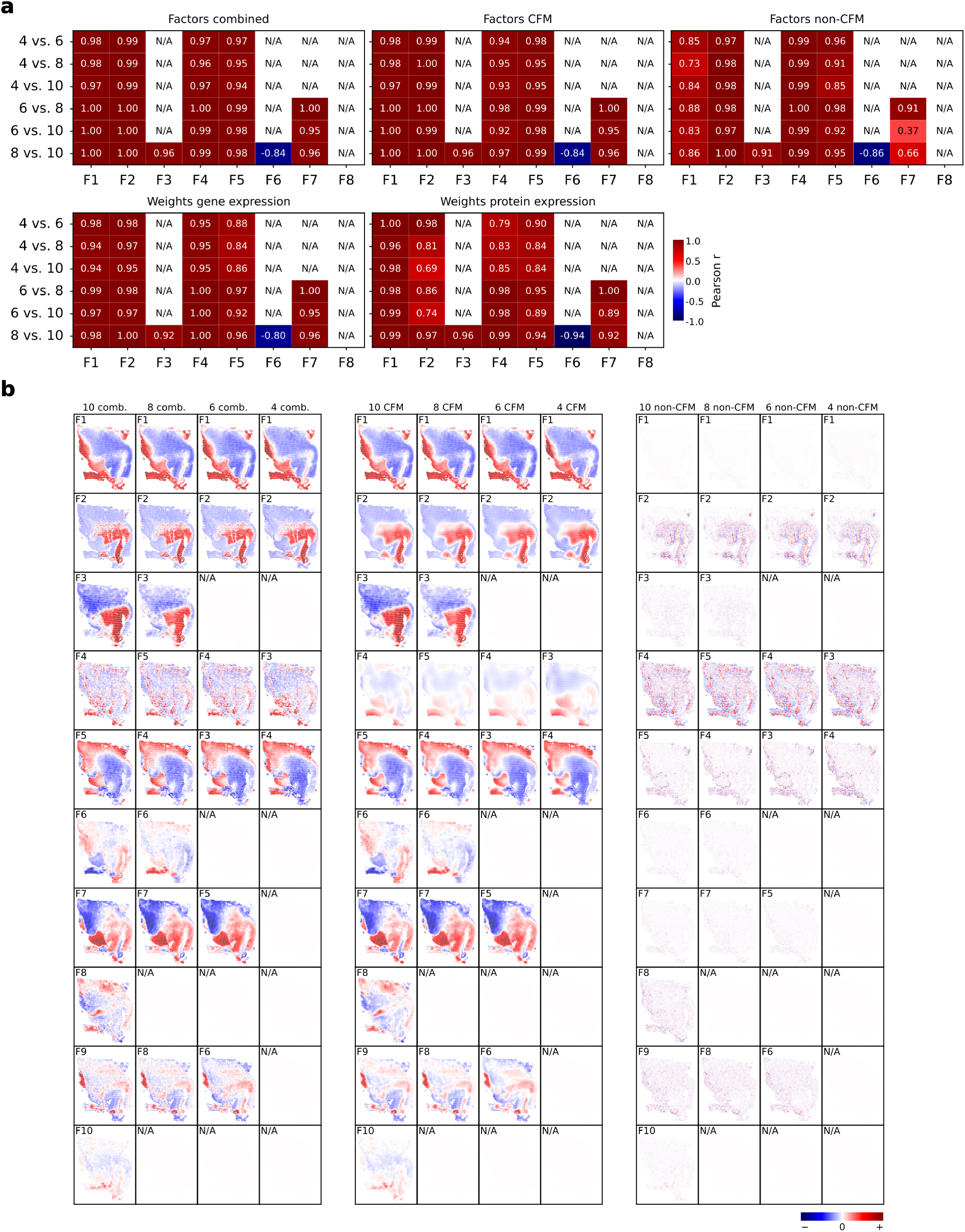
Robustness of MOFTy factors in the glioblastoma dataset across different initializations. **a**, We matched the factors of the eight-, six-, and four-factor configurations to the factors of the ten-factor configuration using linear sum assignment with absolute Pearson correlation as the cost function. **b**, Shown are the Pearson correlation coefficients between the matched factors of the ten-factor configuration and the factors of the eight-, six-, and four-factor configurations. **c**, Shown are the MOFTy factor values over the spatial spots for the ten-factor configuration and the matched factors of the eight-, six-, and four-factor configurations.

**Extended Data Fig. 13.**
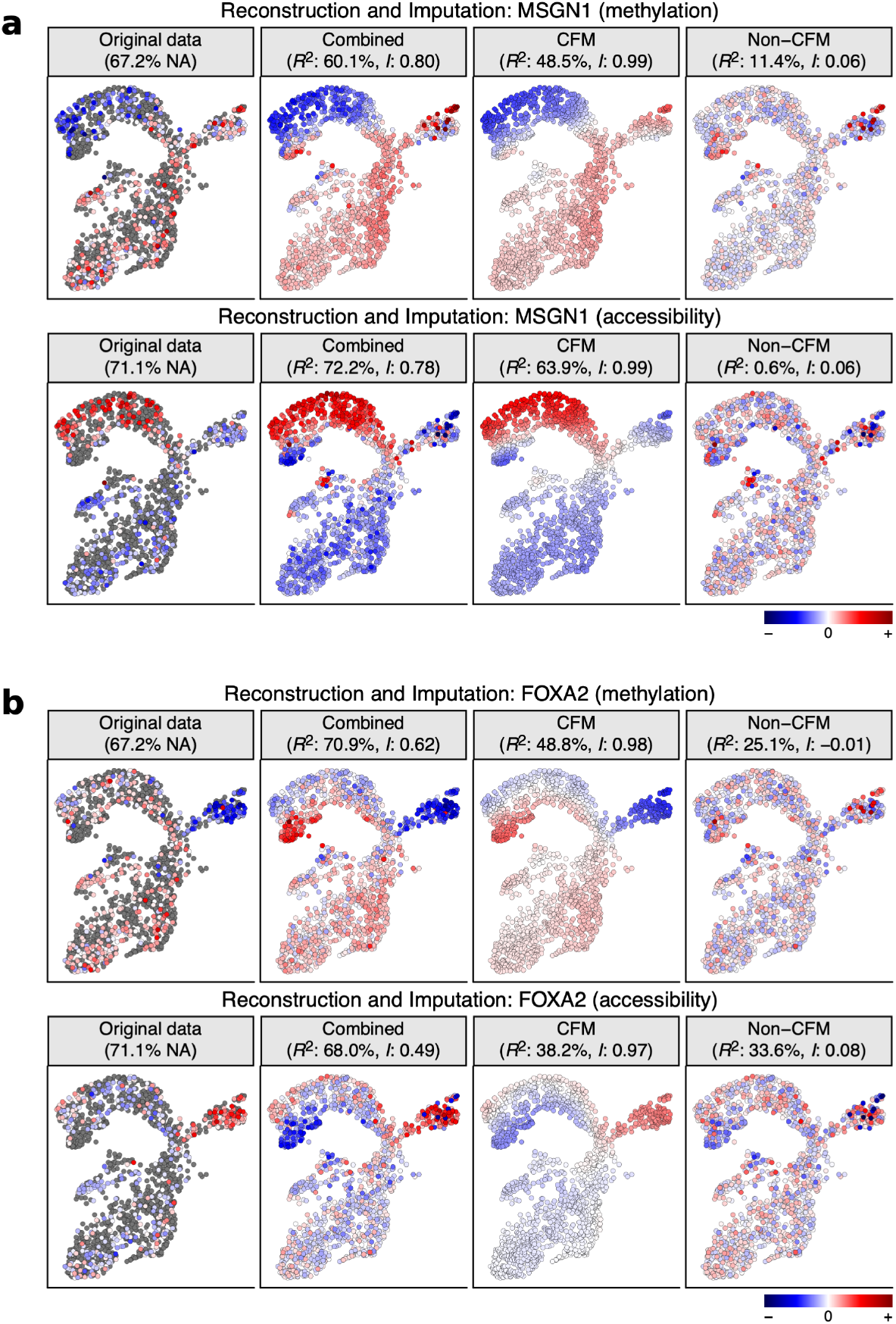
Original observed data (with percentages of missing values) and MOFTy-based reconstructions with missing-value imputation, obtained by multiplying the feature weight vector by the latent factor matrix for the combined, CFM, and non-CFM components (MSGN1, **a**; FOXA2, **b**). Variance explained (*R*^2^) and Moran’s *I* are shown. Each row displays the corresponding values over UMAP coordinates using a shared normalized color scale.

**Extended Data Fig. 14.**
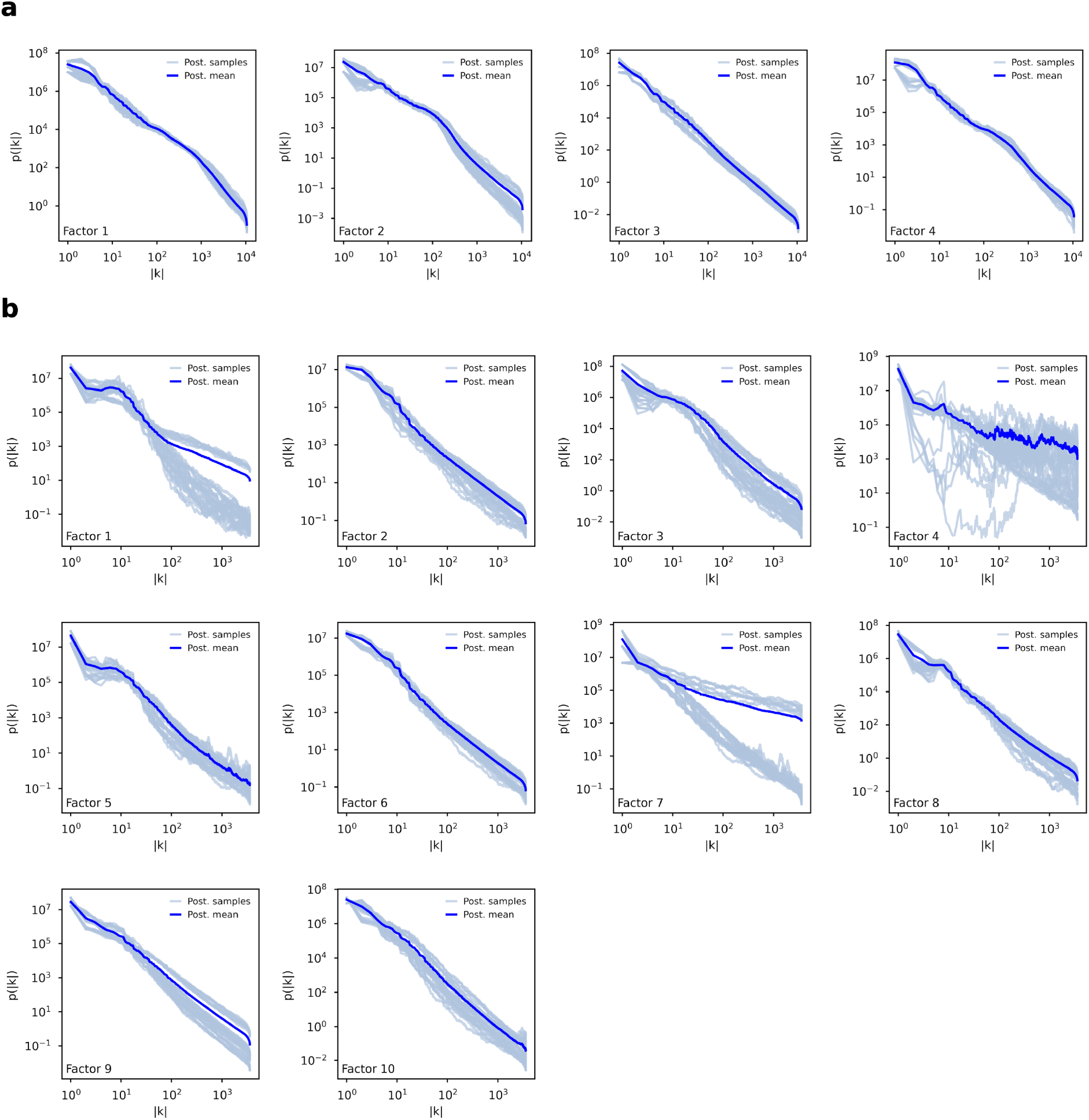
Posterior approximations of power spectra across factors for the glioblastoma (**a**) and mouse gastrulation datasets (**b**). Plotted are the diagonal approximations of the aligned posterior samples (Methods, Section 4.5) and their mean (Post. mean).

## Footnotes

1 For multi-group settings, MEFISTO and MOFA-FLEX combine the covariate-domain kernel with a learned, factor-specific group covariance through a separable product kernel. For a fixed pair of groups, the covariance is therefore given by the covariate-domain kernel multiplied by the corresponding group-pair coefficient. The group covariance modulates the magnitude and sign of dependence across groups but does not alter the functional form of the covariate-domain kernel or introduce additional correlation scales along the covariate. Thus, in the absence of covariate warping, the covariate-domain component remains stationary and isotropic, with a single learned correlation length per factor. When groups share a common covariate grid, the resulting covariance matrix has a Kronecker-product structure. Multi-group settings are not currently supported in MOFTy, and all datasets analyzed in this article comprise a single group.

