## Supplementary Information for "MOFTy: Multimodal Gaussian Process Factor Analysis with Numerical Information Field Theory"

Maximilian Neumann[1,2*], Philipp Arras[3], Anne-Kristin Kaster[2,4], Andreas Ott[1*]

[1]Institute for Mathematics, Heidelberg University, Mathematikon, Im Neuenheimer Feld 205, 69120 Heidelberg, Germany.
[2]Institute for Biological Interfaces 5 (IBG-5), Biotechnology and Microbial Genetics, Karlsruhe Institute of Technology (KIT), Hermann-von-Helmholtz-Platz 1, 76344 Eggenstein-Leopoldshafen, Germany.
[3]No affiliation, Eisenhartstraße 19, 81245 München, Germany.
[4]Institute of Applied Biosciences (IAB), Karlsruhe Institute of Technology (KIT), Fritz-Haber-Weg 4, 76131 Karlsruhe, Germany.

### Overview

The document contains the following supplementary figures:

- Supplementary Figure 1: Synthetic data 1D (900 points).
- Supplementary Figures 2–4: Synthetic data 2D ($30 \times 30$ pixels).
- Supplementary Figures 5–7: Synthetic data 2D ($100 \times 100$ pixels).
- Supplementary Figures 8 and 9: Glioblastoma dataset (four factors).
- Supplementary Figures 10–13: Mouse gastrulation dataset.

For the posterior uncertainty estimates, we show the pointwise or pixelwise posterior uncertainties computed from 32 posterior samples that were aligned and pooled across four seeds, thereby capturing both variational posterior uncertainty and between-seed variability. Cross-correlations between model parameters and higher-order features of the generally non-Gaussian posterior distributions are not reported.

For the 2D synthetic data, we also visualized slices of the posterior samples and posterior means of the combined latent factors across the regular spatial grid points, providing a more detailed illustration of uncertainty in the reconstruction of the latent factors and their components.

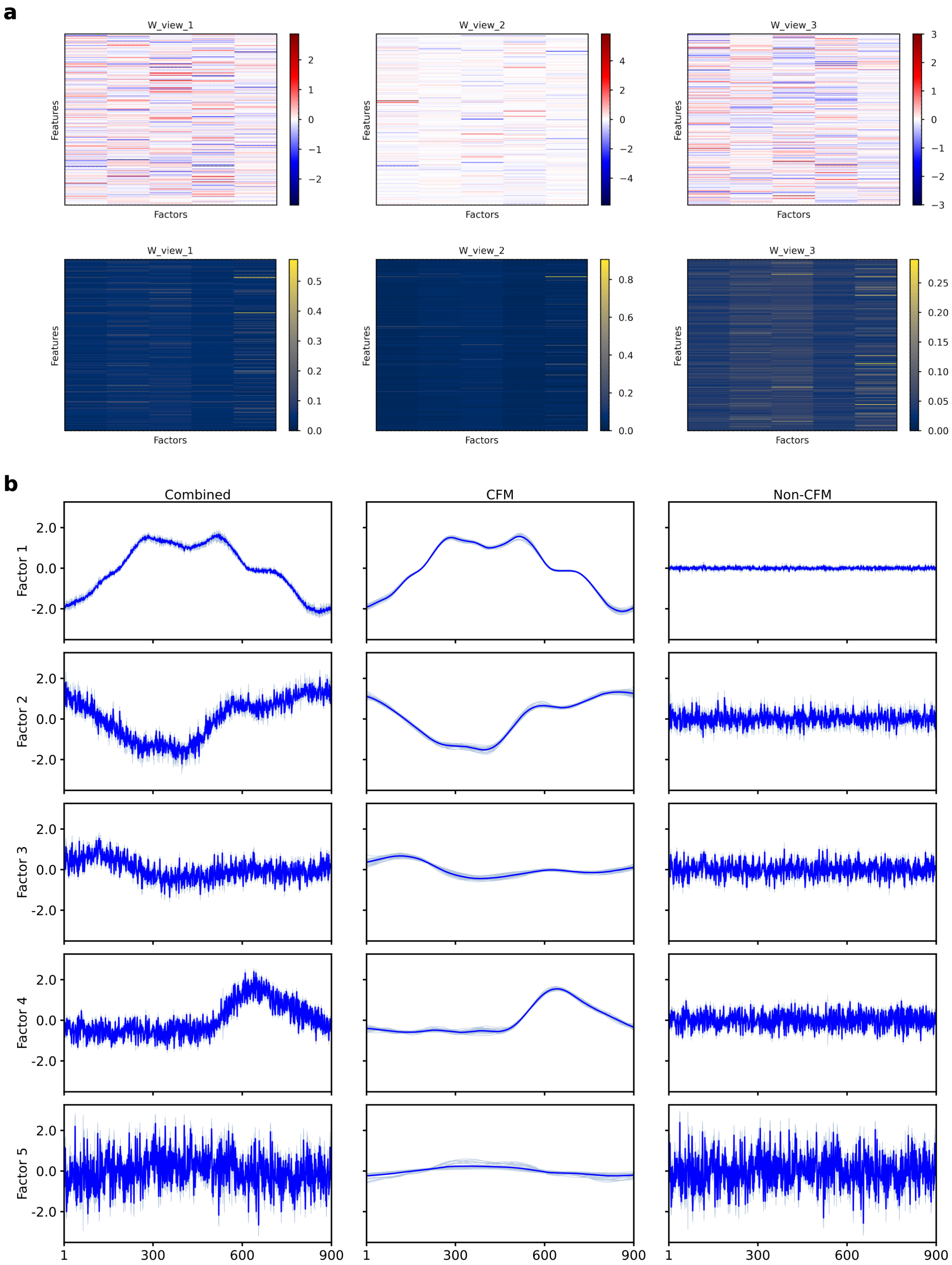


**Fig. 1** Reconstruction and uncertainty for synthetic one-dimensional data (900 points). **a,** The posterior mean (top) and posterior standard deviation (bottom) are shown for the factor weights. **b,** The posterior mean (top) and posterior standard deviation (bottom) are shown for the combined factors and their CFM and non-CFM components. Factors 1–5 are ordered from top to bottom.

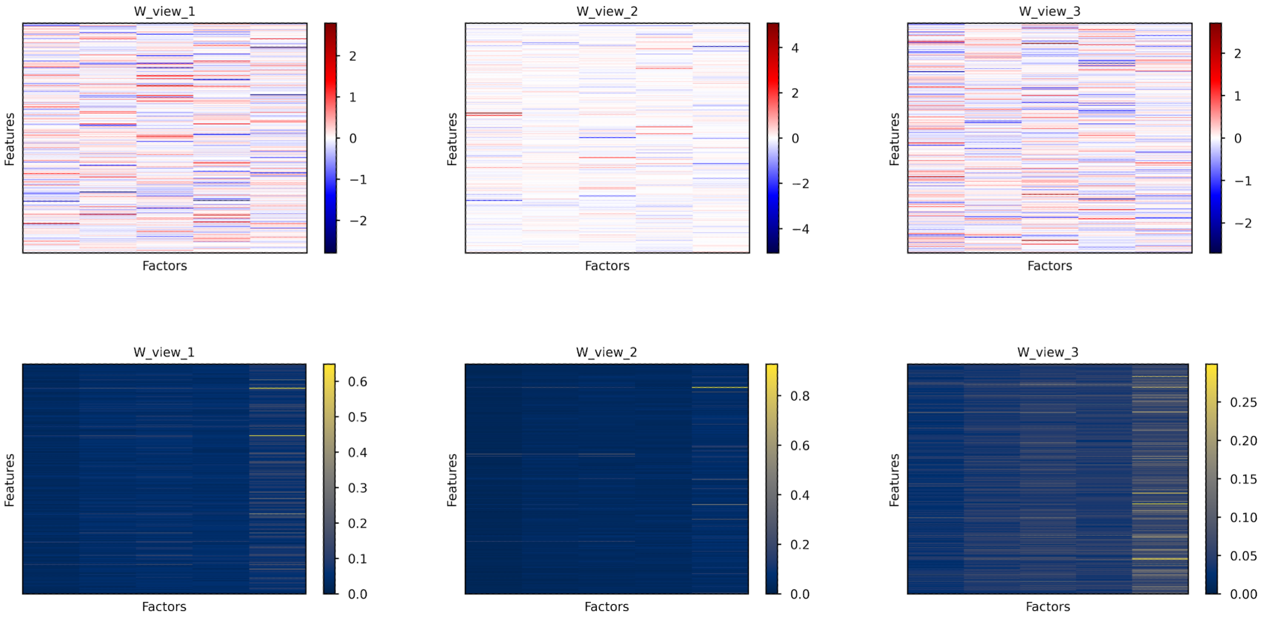


**Fig. 2** Reconstruction and uncertainty for synthetic two-dimensional data (30 × 30 pixels). Shown are the posterior mean (top) and posterior uncertainty, expressed as posterior standard deviation (bottom), of the factor weights.

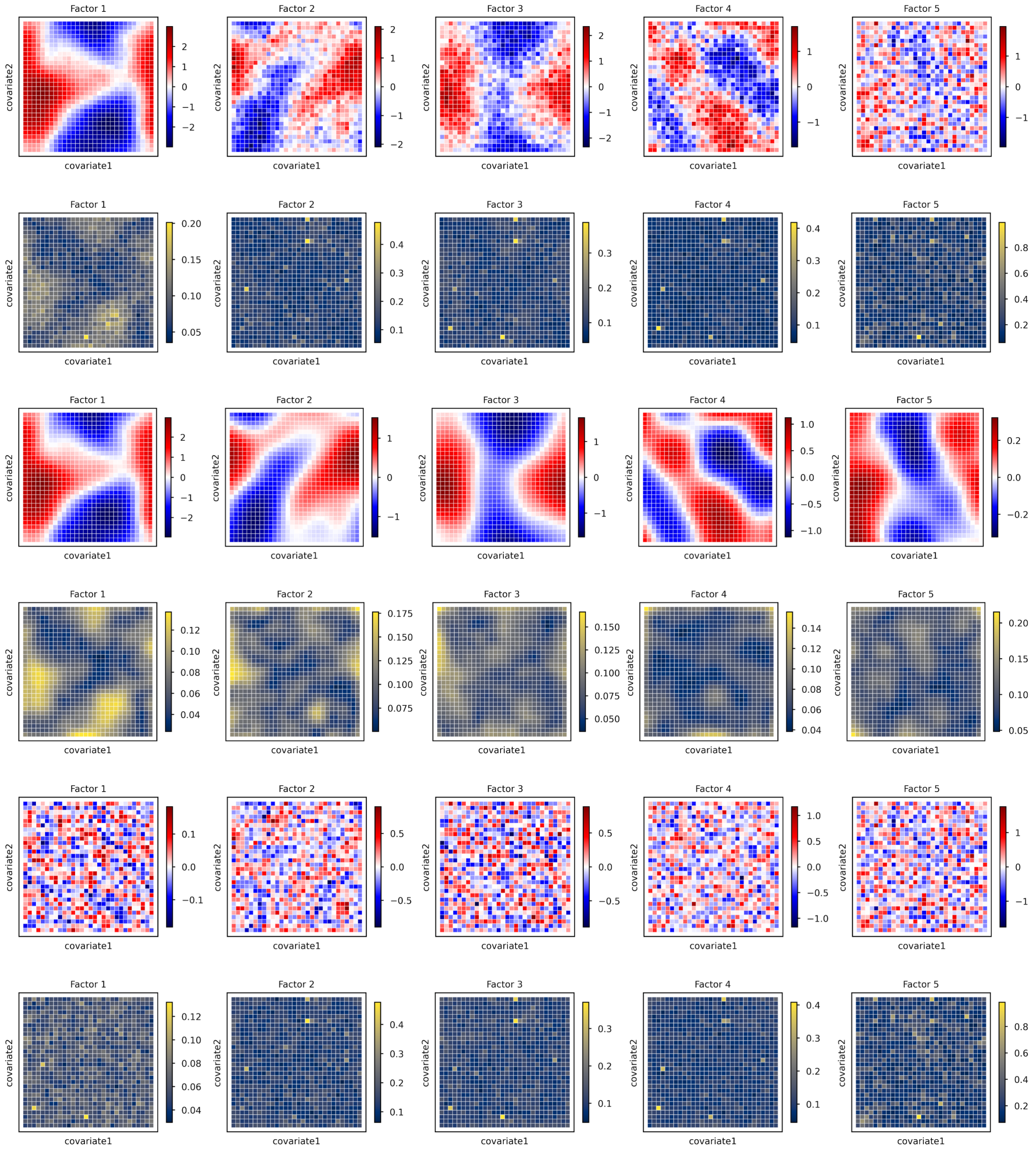


**Fig. 3** Reconstruction and uncertainty for synthetic two-dimensional data (30 × 30 pixels). Rows 1–2 show the posterior mean (top) and posterior uncertainty, expressed as posterior standard deviation (bottom), of the combined factors. Rows 3–4 show the posterior mean (top) and posterior uncertainty, expressed as posterior standard deviation (bottom), of the CFM component. Rows 5–6 show the posterior mean (top) and posterior uncertainty, expressed as posterior standard deviation (bottom), of the non-CFM component.

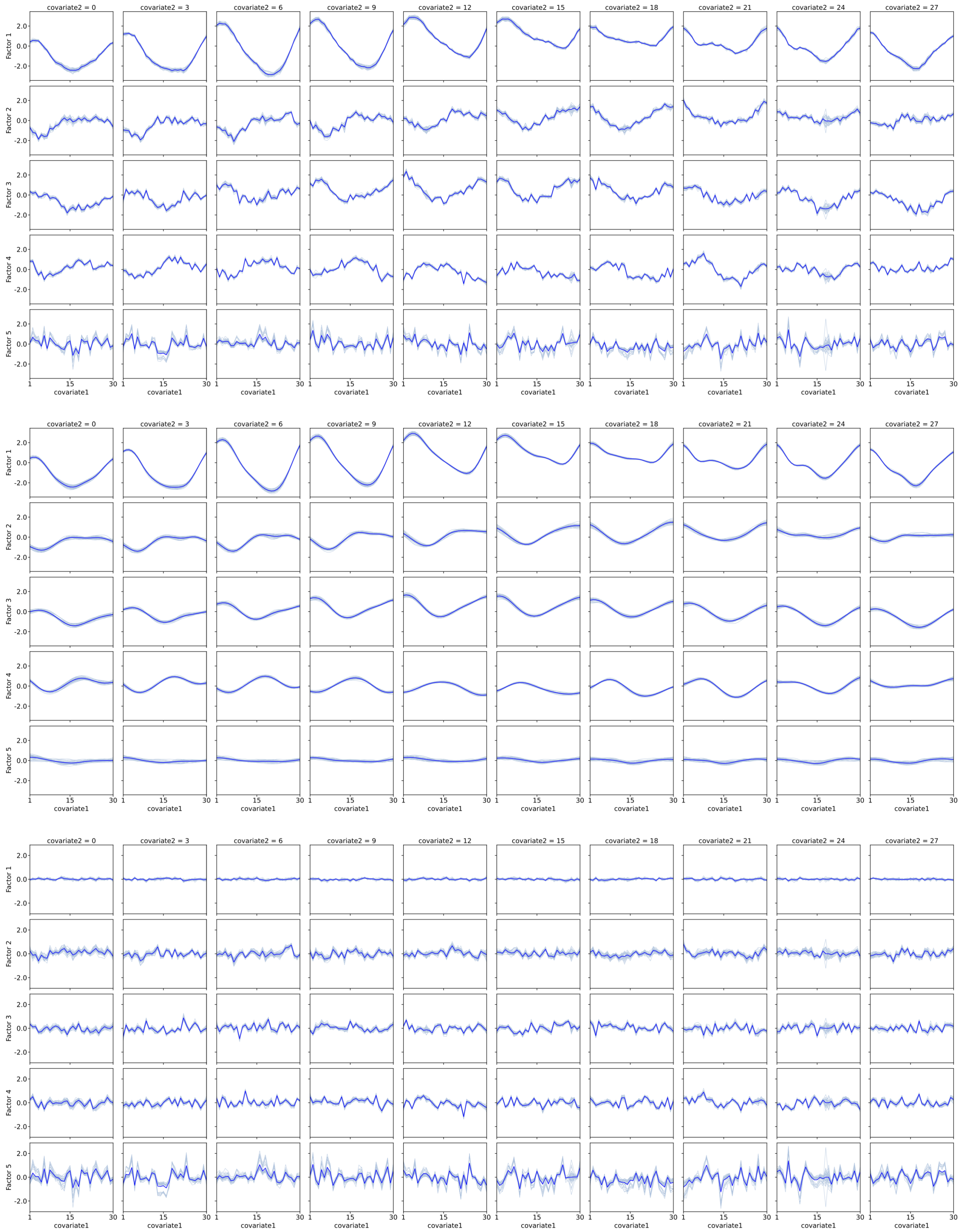


**Fig. 4** Posterior samples (light blue) and posterior means (blue) of the combined (top), CFM (middle), and non-CFM (bottom) factors are shown for 10 slices of the synthetic two-dimensional data ($30 \times 30$ pixels) reconstruction.

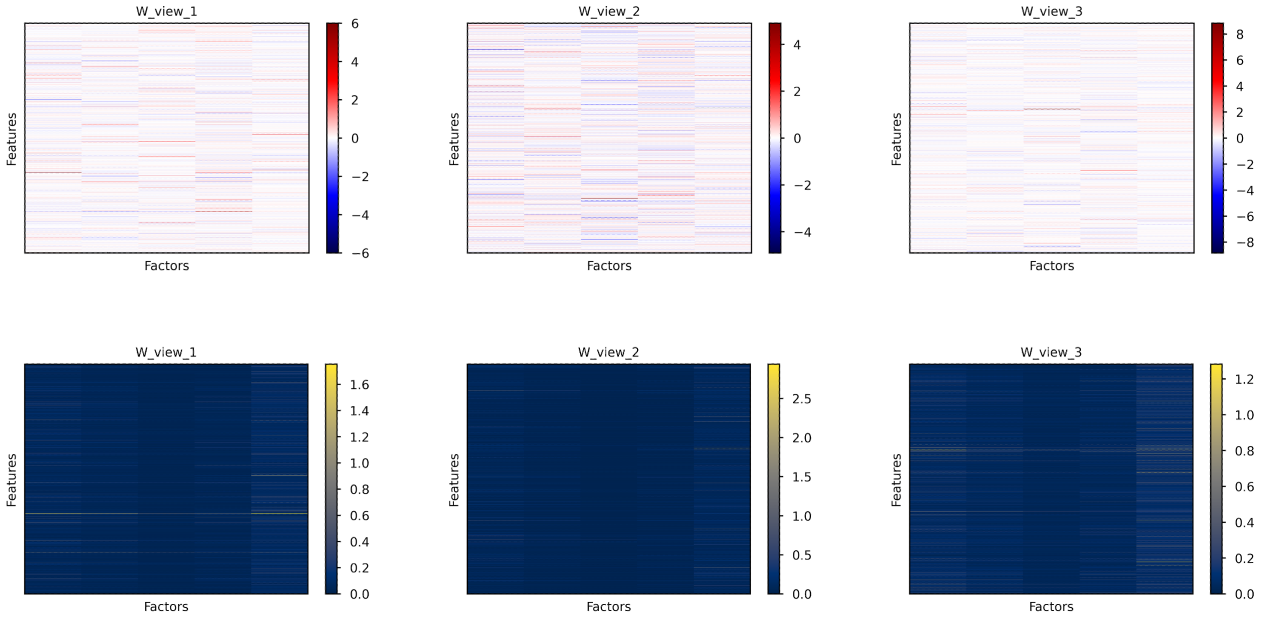


**Fig. 5** Reconstruction and uncertainty for synthetic two-dimensional data ($100 \times 100$ pixels). Shown are the posterior mean (top) and posterior uncertainty, expressed as posterior standard deviation (bottom), of the factor weights.

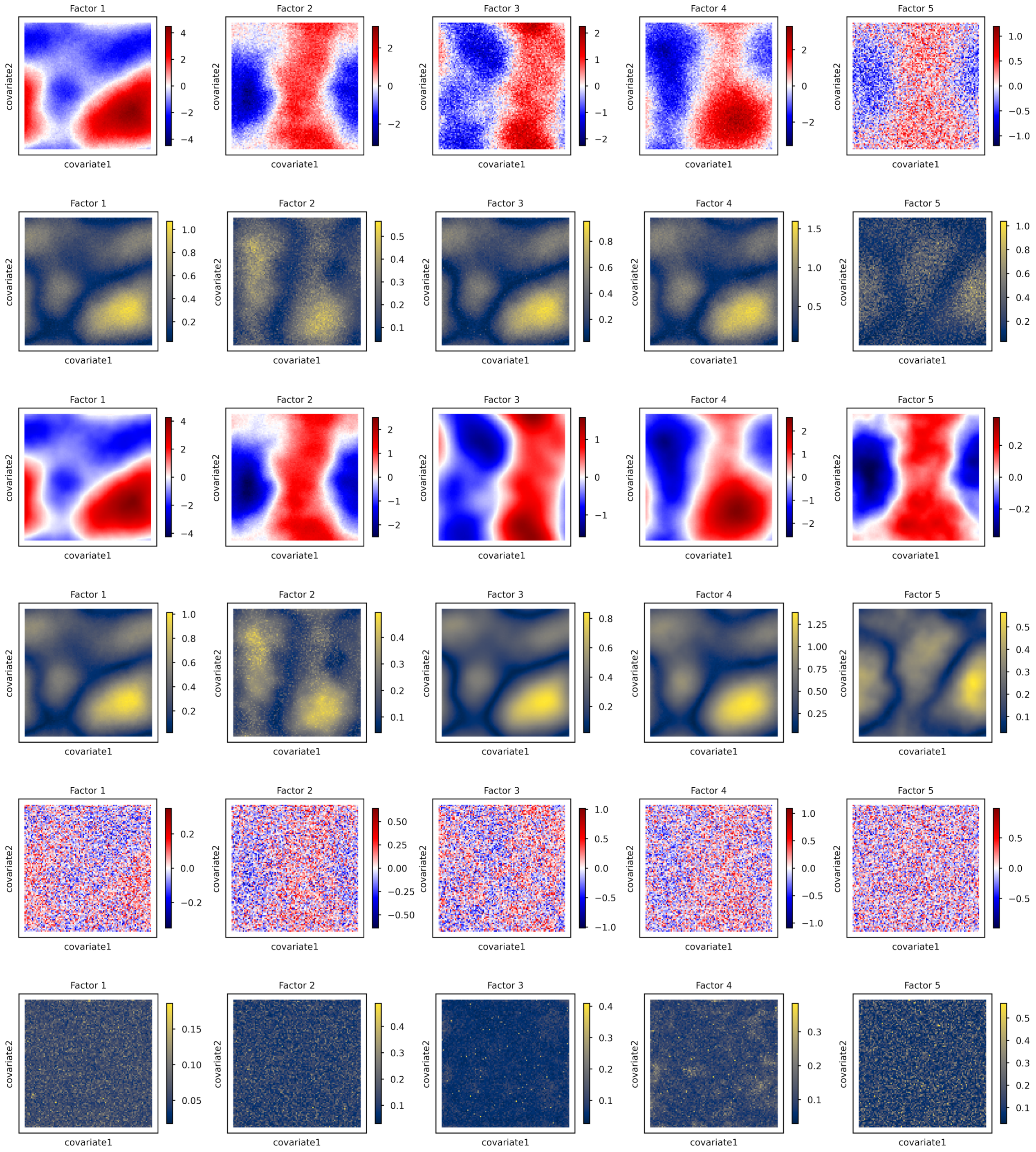


**Fig. 6** Reconstruction and uncertainty for synthetic two-dimensional data (100 × 100 pixels). Rows 1–2 show the posterior mean (top) and posterior uncertainty, expressed as posterior standard deviation (bottom), of the combined factors. Rows 3–4 show the posterior mean (top) and posterior uncertainty, expressed as posterior standard deviation (bottom), of the CFM component. Rows 5–6 show the posterior mean (top) and posterior uncertainty, expressed as posterior standard deviation (bottom), of the non-CFM component.

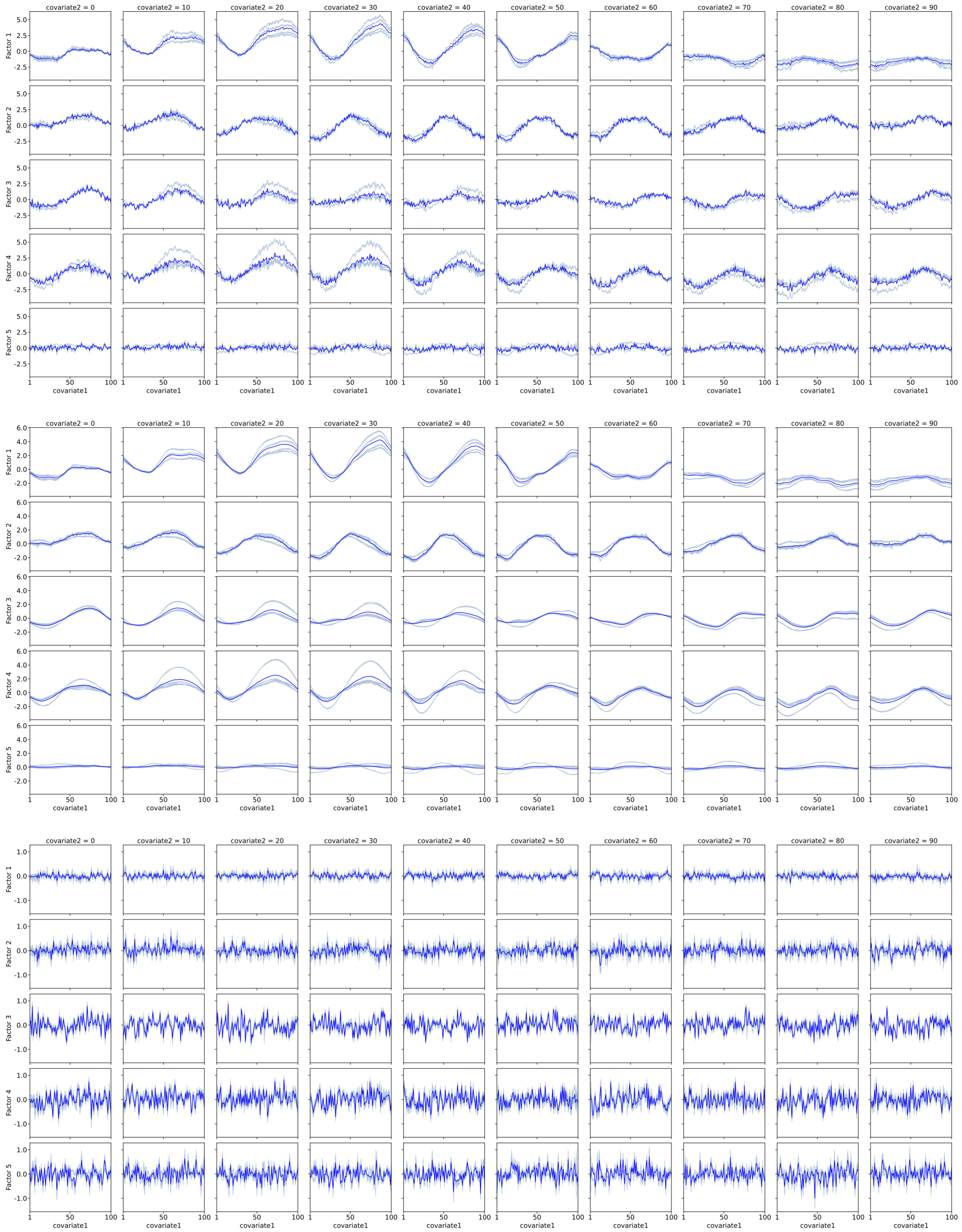


**Fig. 7** Posterior samples (light blue) and posterior means (blue) of the combined (top), CFM (middle), and non-CFM (bottom) factors are shown for 10 slices of the synthetic two-dimensional data (100 × 100 pixels) reconstruction.

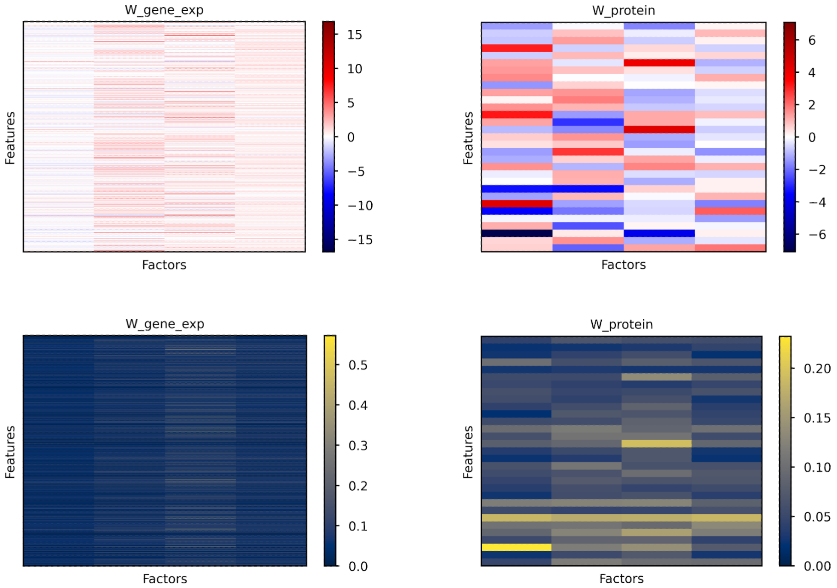


**Fig. 8** Glioblastoma dataset (factor weights). Shown are the posterior mean (top) and posterior uncertainty, expressed as posterior standard deviation (bottom), of the factor weights.

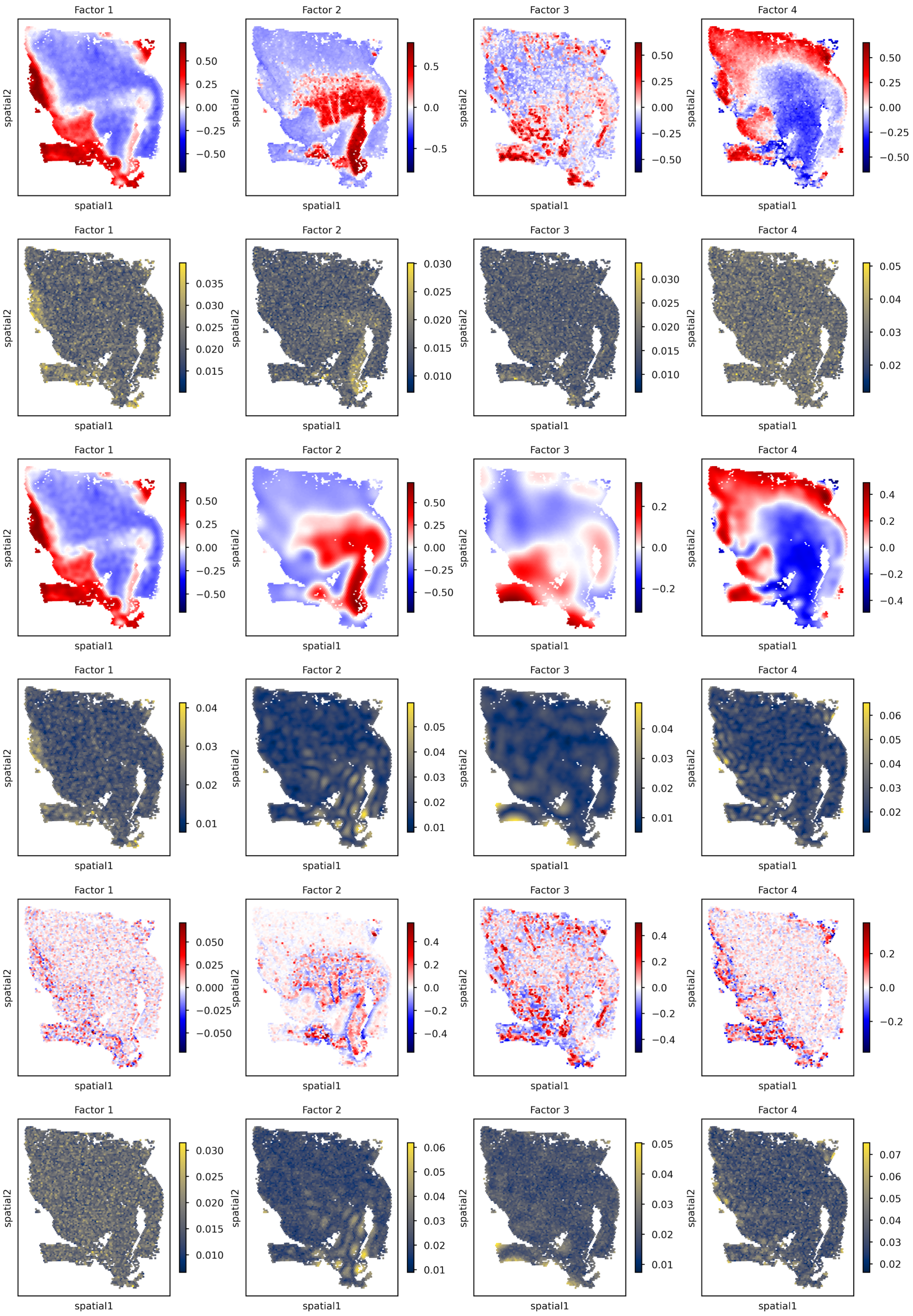


**Fig. 9** Glioblastoma dataset (factors). Rows 1–2 show the posterior mean (top) and posterior uncertainty, expressed as posterior standard deviation (bottom), of the combined factors. Rows 3–4 show the posterior mean (top) and posterior uncertainty, expressed as posterior standard deviation (bottom), of the CFM component. Rows 5–6 show the posterior mean (top) and posterior uncertainty, expressed as posterior standard deviation (bottom), of the non-CFM component.

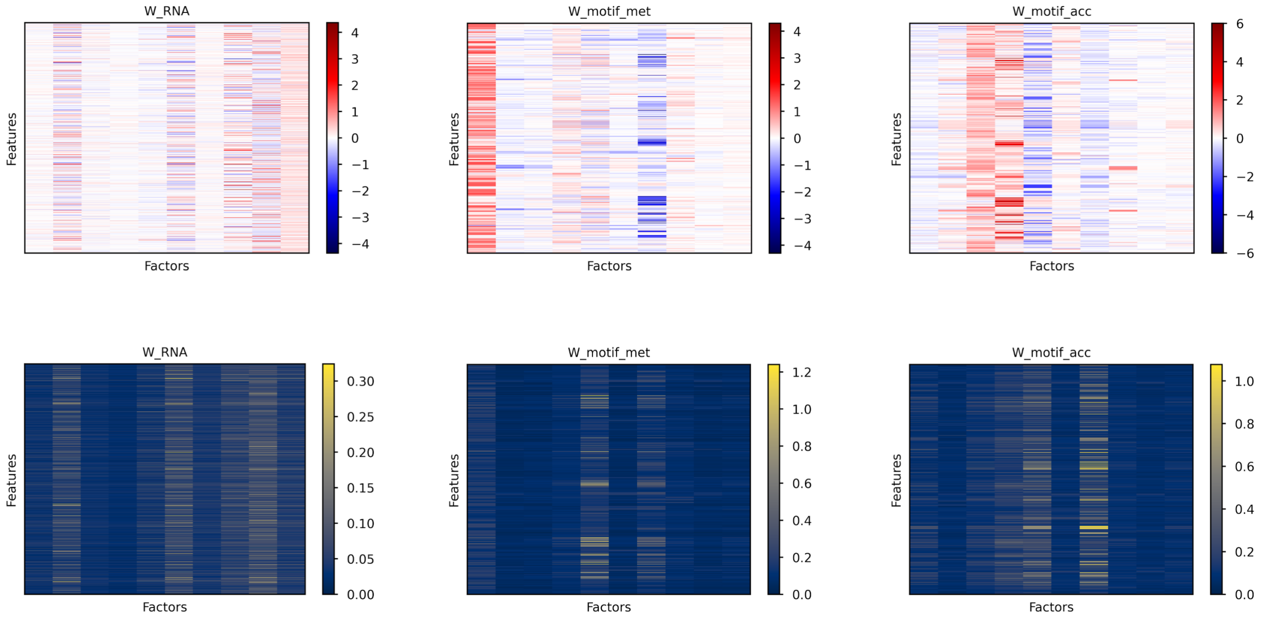


**Fig. 10** Mouse gastrulation dataset (factor weights). Shown are the posterior mean (top) and posterior uncertainty, expressed as posterior standard deviation (bottom), of the factor weights.

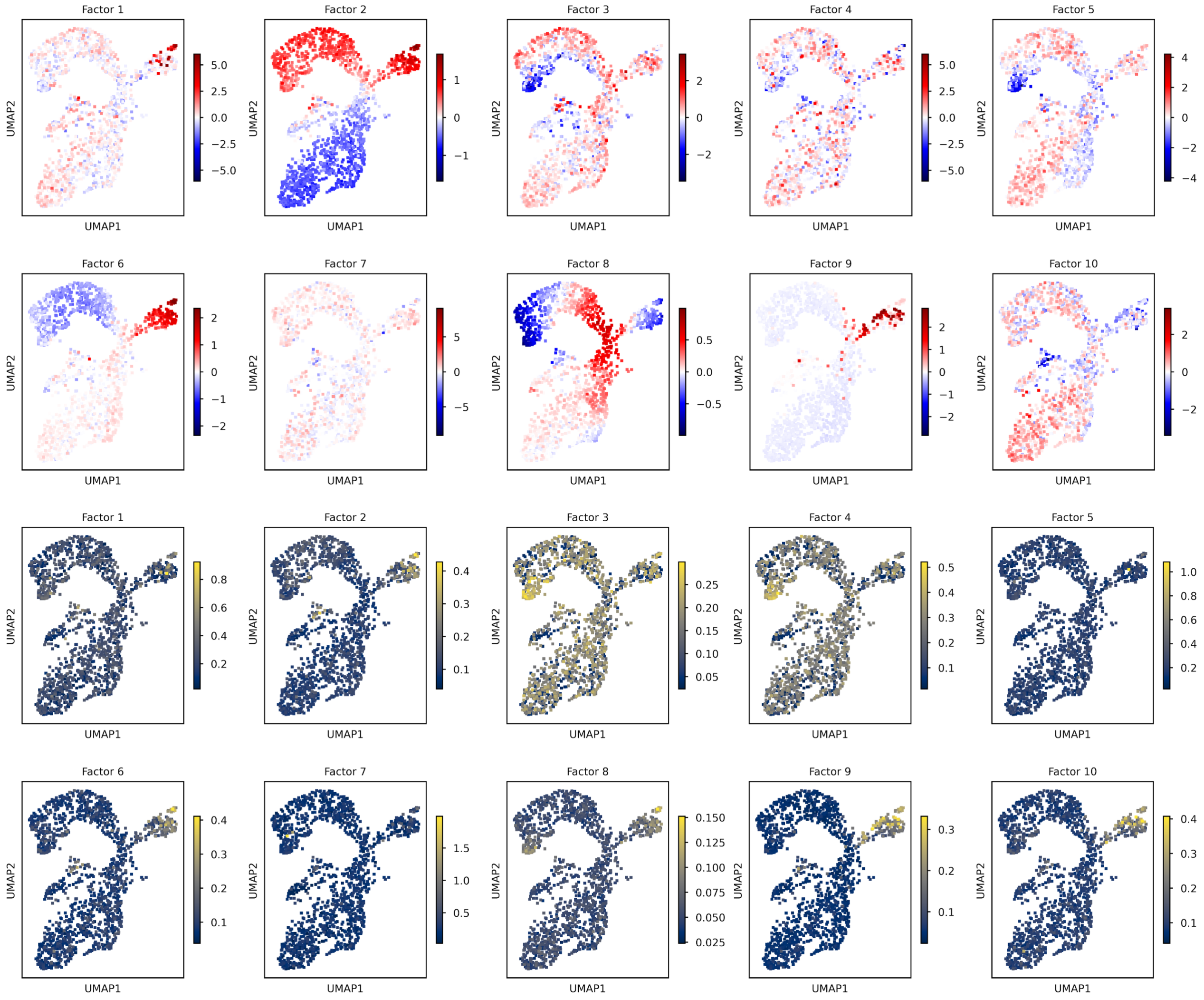


**Fig. 11** Mouse gastrulation dataset (combined factors). Shown are the posterior mean (top) and posterior uncertainty, expressed as posterior standard deviation (bottom), of the combined factors.

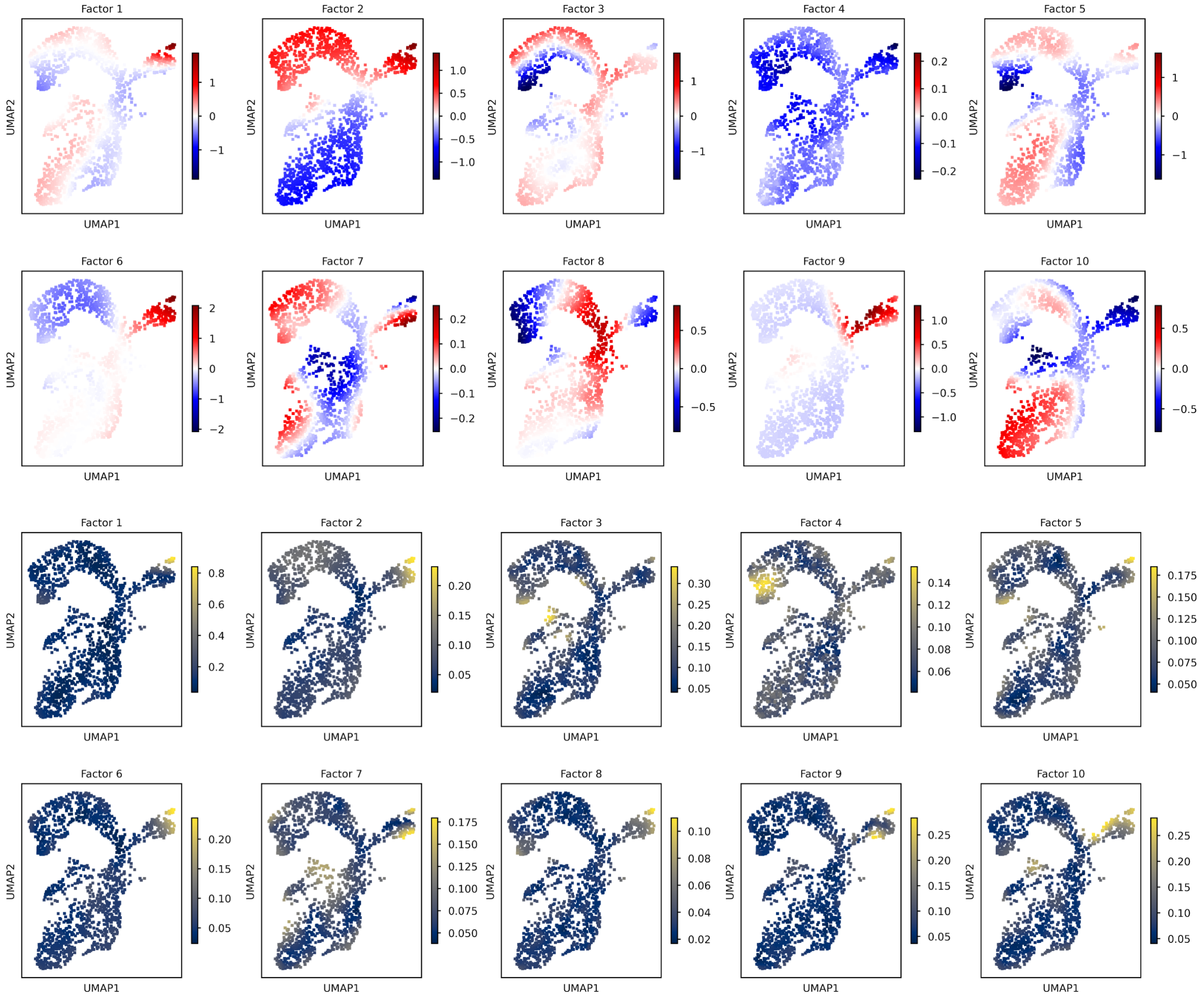


**Fig. 12** Mouse gastrulation dataset (CFM component). Shown are the posterior mean (top) and posterior uncertainty, expressed as posterior standard deviation (bottom), of the CFM component.

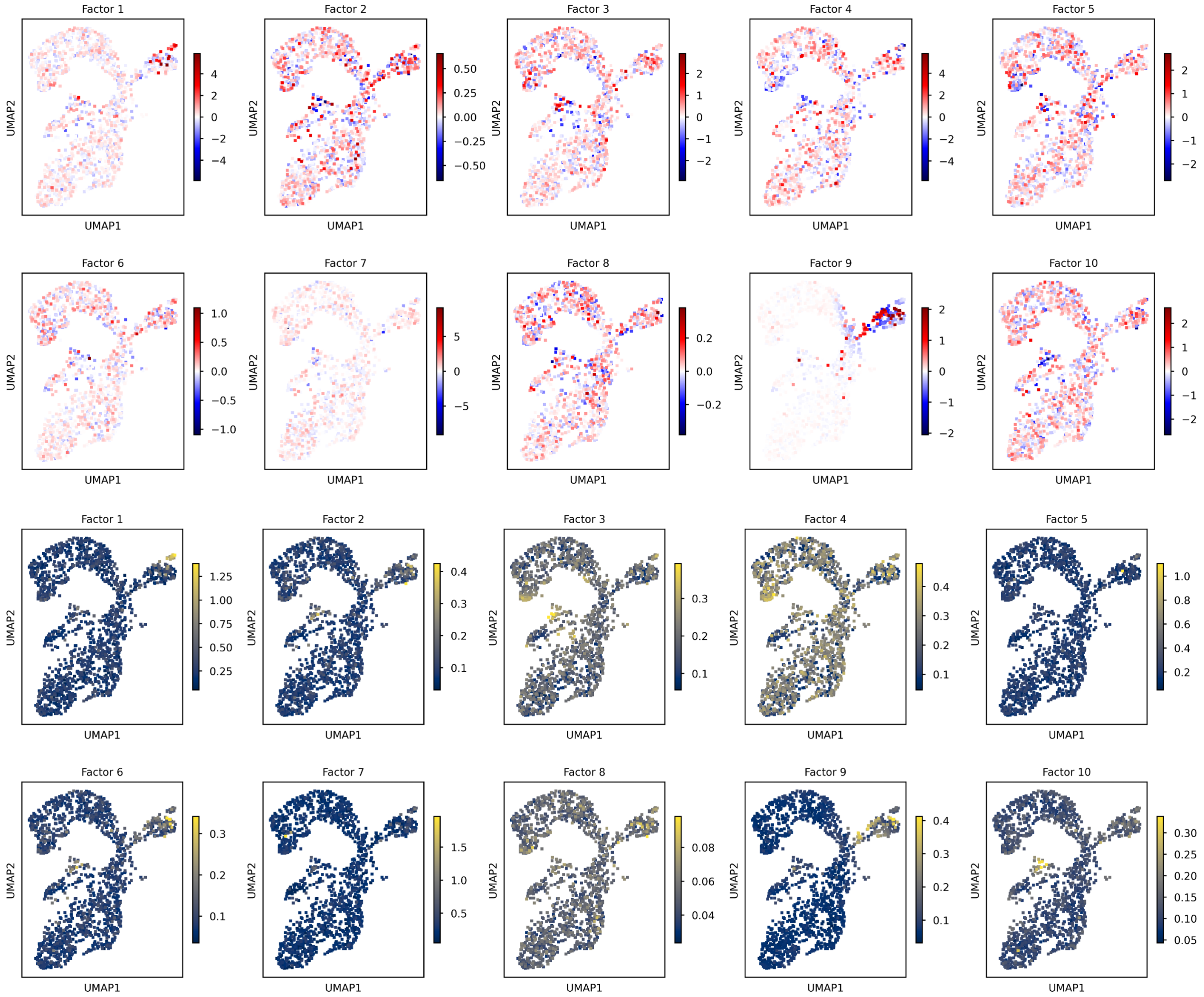


**Fig. 13** Mouse gastrulation dataset (non-CFM component). Shown are the posterior mean (top) and posterior uncertainty, expressed as posterior standard deviation (bottom), of the non-CFM component.
